# A unified computational model for cortical post-synaptic plasticity

**DOI:** 10.1101/2020.01.27.921254

**Authors:** Tuomo Mäki-Marttunen, Nicolangelo Iannella, Andrew G. Edwards, Gaute T. Einevoll, Kim T. Blackwell

## Abstract

Cortical synapses possess a machinery of signalling pathways that leads to various modes of post-synaptic plasticity. Such pathways have been examined to a great detail separately in many types of experimental studies. However, a unified picture on how multiple biochemical pathways collectively shape the observed synaptic plasticity in the neocortex is missing. Here, we built a biochemically detailed model of post-synaptic plasticity that includes the major signalling cascades, namely, CaMKII, PKA, and PKC pathways which, upon activation by Ca^2+^, lead to synaptic potentiation or depression. We adjusted model components from existing models of intracellular signalling into a single-compartment simulation framework. Furthermore, we propose a statistical model for the prevalence of different types of membrane-bound AMPA-receptor tetramers consisting of GluR1 and GluR2 subunits in proportions suggested by the biochemical signalling model, which permits the estimation of the AMPA-receptor-mediated maximal synaptic conductance. We show that our model can reproduce neuromodulator-gated spike-timing-dependent plasticity as observed in the visual cortex. Moreover, we demonstrate that our model can be fit to data from many cortical areas and that the resulting model parameters reflect the involvement of the pathways pinpointed by the underlying experimental studies. Our model explains the dependence of different forms of plasticity on the availability of different proteins and can be used for the study of mental disorder-associated impairments of cortical plasticity.

**Significance statement:** Neocortical synaptic plasticity has been studied experimentally in a number of cortical areas, showing how interactions between neuromodulators and post-synaptic proteins shape the outcome of the plasticity. On the other hand, non-detailed computational models of long-term plasticity, such as Hebbian rules of synaptic potentiation and depression, have been widely used in modelling of neocortical circuits. In this work, we bridge the gap between these two branches of neuroscience by building a detailed model of post-synaptic plasticity that can reproduce observations on cortical plasticity and provide biochemical meaning to the simple rules of plasticity. Our model can be used for predicting the effects of chemical or genetic manipulations of various intracellular signalling proteins on induction of plasticity in health and disease.

## 1 Introduction

Synaptic plasticity in the neocortex has been under intense research since the first observations of neocortical long-term potentiation (LTP) [Komatsu et al., 1981, Lee, 1982]. Although most often studied in brain slices, synaptic plasticity in the neocortex is a key phenomenon underlying vital mammalian brain processes ranging from formation and storage of memories to attentional selection [Roelfsema and Holtmaat, 2018]. These processes are impaired in heritable mental illnesses such as schizophrenia and fragile X syndrome, as well as neurodegenerative diseases such as Alzheimer’s disease, all of which have been associated with deficits in cortical plasticity [Kantrowitz et al., 2017, Martin and Huntsman, 2012, Koch et al., 2014]. Improved understanding of neocortical synaptic plasticity all the way from molecular to circuit level is therefore needed to further our understanding of these yet incurable diseases.

Similar to hippocampal synaptic plasticity [Larkman and Jack, 1995], synaptic plasticity in the neocortex is highly variable — the outcomes of any plasticity-inducing protocol depends on the cortical area, neuron type as well as details of the stimulation protocol [Castro-Alamancos et al., 1995, Froc and Racine, 2005, Sjostrom et al., 2008, Feldman, 2009]. Computational models provide a tool for efficient hypothesis testing of mechanisms of neocortical plasticity, which helps to overcome the challenges posed by excessive variability. The foundations of our mechanistic understanding of neocortical synaptic plasticity lie upon the phenomenological Bienenstock-Cooper-Munro (BCM) theory, which predicts that small synaptic Ca^2+^ transients cause long-term depression (LTD) whereas large Ca^2+^ transients give rise to LTP [Bienenstock et al., 1982]. Simple BCM-based models and the closely related models of spike-timing-dependent plasticity (STDP) have been widely used to explain the emergence of input-specific cell assemblies mediating, e.g., orientation selectivity [Shouval et al., 1997] or memory traces [Klampfl and Maass, 2013] in the cortex. These models, however, typically fail to provide a mechanistic understanding of the biochemistry within the synapse — namely, they do not reveal how various molecules downstream of Ca^2+^ regulate the induction and maintenance of plasticity occurring in neuronal circuits, their composite neurons and synapses of the cortex. Moreover, current models often ignore the joint contributions of neuromodulators, which are critical for inducing some forms of cortical synaptic plasticity [Meunier et al., 2017, Brzosko et al., 2019]. These shortcomings impede testing biochemical mechanisms of heritable mental illnesses associated with impaired cortical plasticity.

In this work, we aim at filling this gap of knowledge by introducing a biochemically detailed, mass-action law-based model of neocortical post-synaptic plasticity that can be used to study the induction of plasticity in different genetic conditions and neuromodulatory states, and under various stimulation protocols. Despite the lack of biochemically detailed models of synaptic plasticity in the neocortex, models of intracellular signalling have been used to study LTP and LTD in the hippocampus [Bhalla and Iyengar, 1999, Jedrzejewska-Szmek et al., 2017], cerebellum [Gallimore et al., 2018], and striatum [Blackwell et al., 2018]. These models permit systematic studies on how patterns of Ca^2+^ inputs to the post-synaptic spine, either alone or in combination with neuromodulatory actions, activate different signalling pathways leading to post-synaptic plasticity in the form of, e.g., AMPA-receptor (AMPAR) phosphorylation and membrane insertion. We integrate quantitative descriptions of the intracellular signalling pathways underlying synaptic plasticity in the neocortex into a unified model that is capable of describing both stimulation protocol-dependent plasticity, as well as neocortically observed neuromodulator-gated forms of STDP. We show that our model can be tuned by alterations of protein expression to reproduce not only BCM-like forms of plasticity but also experimental observations on neocortical plasticity from various cortical areas. Our results help to quantify and explain the differences in molecular constituents of different forms of neocortical LTP and LTD, and the different, data-fitted versions of our model can be directly used to examine the effects of chemical inhibitors and genetic manipulations of signalling proteins on synaptic plasticity in different cortical cells.

## 2 Methods

### 2.1 Construction and calibration of the biochemically detailed model of post-synaptic plasticity in the cortex

We created a model of pathways leading from Ca^2+^ inputs and activation of *β*-adrenergic receptors, metabotropic glutamate receptors, and muscarinic acetylcholine receptors to the phosphorylation and insertion of AMPARs into the membrane. We started by using the model of [Jedrzejewska-Szmek et al., 2017] for GluR1 phosphorylation at sites S831 and S845, which are phosphorylated by protein kinase A (PKA) and Ca^2+^/calmodulin-dependent kinase II (CaMKII), respectively, as a basis for our unified model. We added the metabotropic glutamate receptor (mGluR) and muscarinic acetylcholine M1 receptor-dependent pathways leading to protein kinase C (PKC) activation from [Kim et al., 2013] and [Blackwell et al., 2018], respectively, and adopted the PKC-dependent endocytosis of GluR2 and reinsertion to the membrane from [Gallimore et al., 2018] as these pathways are critical for neocortical plasticity [Seol et al., 2007]. As we included molecular species from different models and as we omitted certain molecular species that affected the dynamics of the underlying species but were not imperative for the pathways we wanted to describe, calibration of the model reactions was necessary. Following [Hayer and Bhalla, 2005], we allowed the insertion and removal of GluR1 subunits to and from the membrane that depended on their state of S845 phosphorylation. We also allowed spontaneous membrane insertion of non-S845-phosphorylated GluR1; we chose the rate of this reaction so that on average one fifth of the (non-S845-phosphorylated) GluR1s were membrane-expressed in steady state, as suggested by experimental data [Oh et al., 2005]. We adjusted the forward rate of GluR2 insertion to the membrane to decrease the proportion of membrane-inserted vs. internalised GluR2s (Fig. 1A), following experimental data according to which 45% of GluR2s were membrane-inserted at resting conditions [Ashby et al., 2004]. We also adopted the three-step calmodulin (CaM) activation of [Gallimore et al., 2018] instead of the two-step activation of [Jedrzejewska-Szmek et al., 2017] where the reaction rates of CaM binding two Ca^2+^ ions were linearly dependent on the number of Ca^2+^ ions. The response curve for CaM activation by Ca^2+^ was steeper in this model (Fig. 1B). As a result, our model predicted a more prominent activation of CaM in response to a large influx of Ca^2+^ than the model of [Jedrzejewska-Szmek et al., 2017] (Fig. 1C).

**Figure 1:**
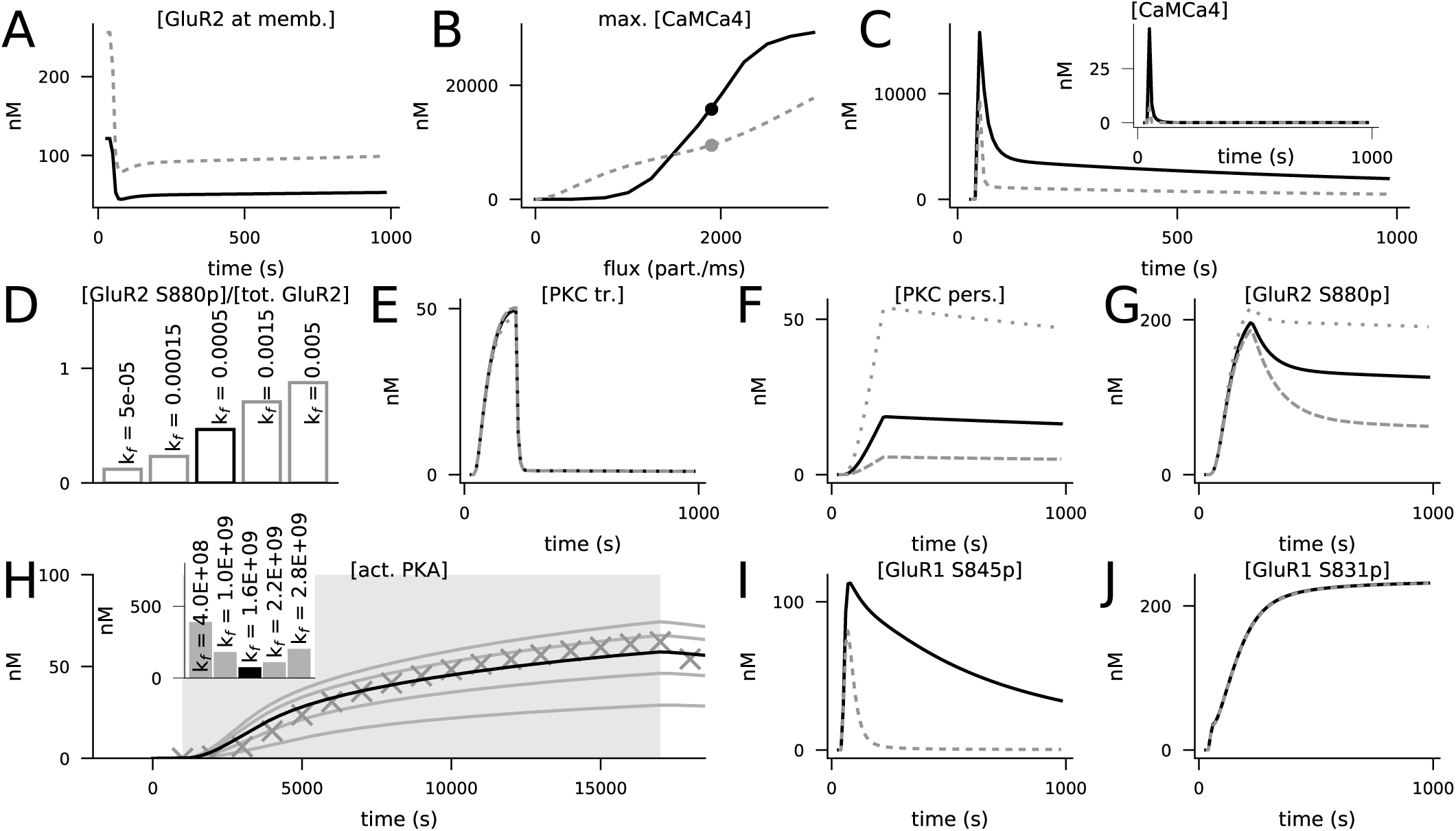
Calibration of the model. Black curves represent the final model, while grey lines represent predictions of models where previous model components or tentative parameter values were used. **A**: Concentration of membrane-inserted GluR2 in 4xHFS when the forward rate of the membrane insertion of non-phosphorylated GluR2 was 0.0055 1/ms [Gallimore et al., 2018] (grey) or 0.00025 1/ms (black). The rate 0.00025 1/ms caused a resting-state concentration of 121 nM for the membrane-bound GluR2 subunits, which is 45% of the total GluR2 concentration (270 nM). **B**: Maximum concentration of total activated CaM in response to 4xHFS as a function of Ca^2+^ input amplitude when the two-step (grey) or three-step (black) activation of CaM by Ca^2+^ was used. The dots represent the data of (C). **C**: Concentration of non-protein-bound activated CaM (inset) or total activated CaM (main figure) in response to 4xHFS when the two-step (grey) or three-step (black) activation of CaM by Ca^2+^ was used. **D**: Percentage of S880-phosphorylated GluR2 15 minutes after LFS when different forward rates of the activation of persistent PKC (*k_f_* between 0.00005 and 0.005 1/(nM ms)) were used. The value *k_f_* = 0.0005 1/(nM ms) gave a percentage of 47%, in close agreement with [Ashby et al., 2004]. **E–G**: The dynamics of transiently active PKC (E) were not strongly influenced by the forward rate of the activation of persistent PKC (reaction 140), but those of persistently active PKC (F) and S880-phosphorylated GluR2 (G) were significantly affected. Black curves show the data corresponding to *k_f_* = 0.0005 1/(nM ms), while the grey lines show the data corresponding to *k_f_* = 0.00015 1/(nM ms) (dashed) and *k_f_* = 0.0015 1/(nM ms) (dotted). **H**: Predicted responses of an isolated PKA activation model (reactions 59 and 93) to a 16-second cAMP input (dim grey background) when different values of the forward rate of PKA binding with four cAMP molecules were used. The curves show the concentration of the catalytic PKA subunit when different forward rates of PKA–cAMP binding (from bottom to top: 0.4*×*10^9^, 1.0*×*10^9^, 1.6*×*10^9^, 2.2*×*10^9^, and 2.8*×*10^9^ 1/(nM^4^ms)) were used. The markers show the corresponding data when the two-step PKA–cAMP binding model of [Jedrzejewska-Szmek et al., 2017] was used. Inset: summed absolute differences between the tentative data (curves) and simulated data from the previous model (markers). The model with the forward rate of *k_f_* = 1.6*×*10^9^ 1/(nM^4^ms) gave the closest correspondence to the model of [Jedrzejewska-Szmek et al., 2017]. **I**: Concentration of S845-phosphorylated GluR1 in response to 4xHFS when the single-step (reaction 59, black) or two-step (from [Jedrzejewska-Szmek et al., 2017], grey) PKA–cAMP binding was used. **J**: Concentration of S831-phosphorylated GluR1 in response to 4xHFS when PKC did (black) or did not (grey, overlaid) phosphorylated S831 in GluR1s.

To allow long-term activation of PKC, we adopted a persistently activated form of PKC, mediated by arachidonic acid (AA), from [Hellgren Kotaleski et al., 2002]. We calibrated the rates of this reaction as follows. The backward rate was chosen so that approximately 90% of PKC would be active after 10 min, inspired by experimental data of [Shirai et al., 1998]. The forward rate was chosen so that low-frequency stimulation (LFS) with effective PLC activation led to approximately 50% of the GluR2s being phosphorylated (Fig. 1D), following experimental data [Ahmadian et al., 2004]. The implications of these adjustments on the dynamics of transiently activated PKC, persistently activated PKC and GluR2 S880 phosphorylation are illustrated in Fig. 1E, 1F, and 1G, respectively.

We adopted the simplified, mass-action law-based PKA activation model (reaction 59; Tab. 1) from [Williamson et al., 2009] (where it was called model “C”) instead of the 2-stage, linearised cAMP-binding of PKA in [Jedrzejewska-Szmek et al., 2017] and [Blackwell et al., 2018]. We fitted the forward rate to data simulated with the original model (Fig. 1H) to produce a longer-lasting S845 phosphorylation (typically, *>*10 min duration of S845 phosphorylation was observed [Seol et al., 2007, Xue et al., 2014]) in the 4xHFS protocol (Fig. 1I). To account for the experimental observation the PKC phosphorylates GluR1 at site S831 [Roche et al., 1996], we added this reaction using the rates identical to those of GluR1-S831 phosphorylation by phosphorylated CaMKII (see reactions 71–72). However, the presence of this reaction did not have a large effect on the S831 phosphorylation of GluR1 under standard conditions (Fig. 1J). Finally, we introduced an immobile Ca^2+^ buffer with a Ca^2+^ binding rate of 0.0004 1/(nM ms), a release rate 20.0 1/ms, and an initial concentration of 500 µM (these values are within the range of experimental observations and values used in models [Matthews and Dietrich, 2015]). The model reactions are described in Tab. 1 and the initial concentrations are listed in Tab. 2.

**Table 1:**
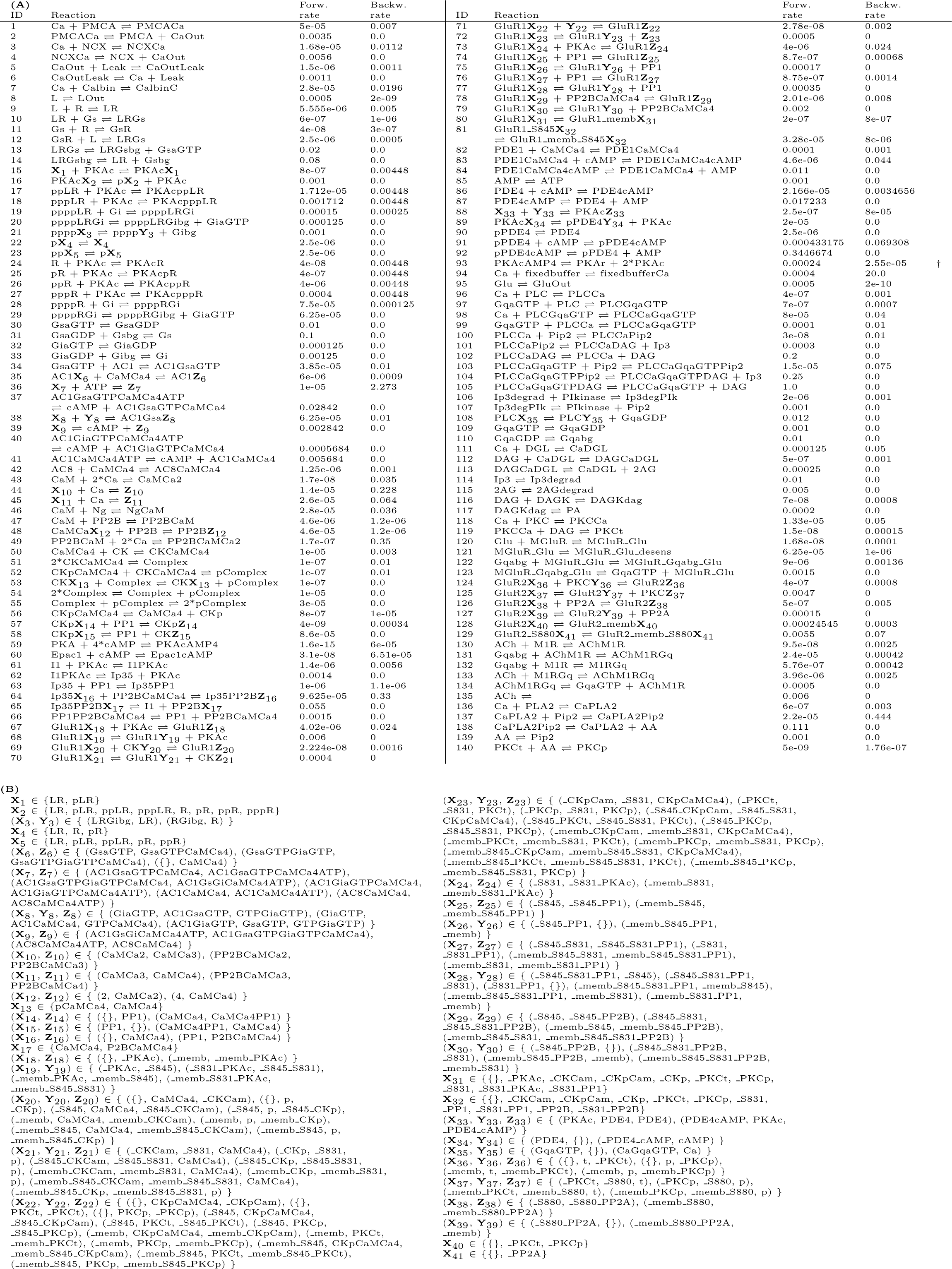
List of model reactions. **A**: The reaction-rate units are in 1/ms, 1/(nMms), 1/(nM^2^ms), 1/(nM^3^ms), or 1/(nM^4^ms), depending on the number of reactants. Reactions are grouped by similar modes of action and identical forward and backward rates. The denominators **X**, **Y**, and **Z** represent groups of species detailed below. *†*: backward reaction rate proportional to [PKAc], not to [PKAc]^2^. **B**: Groups of species as used in panel (A).

**Table 2:**
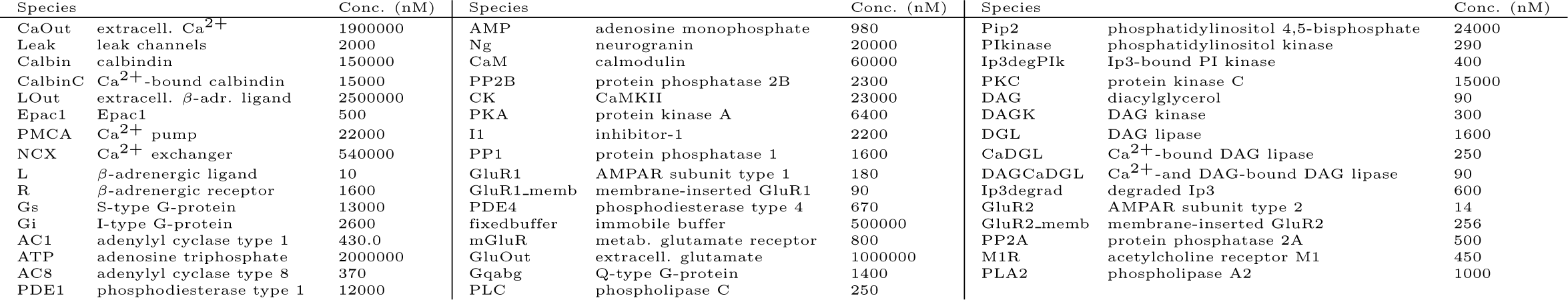
List of initial concentrations of molecular species. All non-mentioned species have an initial concentration of 0 nM.

Throughout the work, we simulated the signalling pathways in a single compartment representing a dendritic spine of size 0.5 µm^3^. In reality, some of the molecular species are prevalently present in the cytosol, some attached to the membrane, some in the extracellular medium in an immediate vicinity to the membrane, and others outside the cell further away from the synaptic cleft (free in the extracellular medium or sequestered to other cells). As commonly done in the field, we solved this problem by introducing species that represent a molecular species confined in a particular location: reactions 1–6 describe the expulsion of Ca^2+^ from the cytosol into the extracellular medium, reactions 8, 95, and 135 describe the escape of ligands from the vicinity of the synapse, and reactions 80–81 and 128–129 for the translocation of the AMPARs to/from the membrane (Tab. 1).

### 2.2 Statistical model for numbers of AMPAR tetramers at the membrane and the total synaptic conductance

AMPARs have different conductances depending on their subunit composition and phosphorylation state [Oh and Derkach, 2005], but it is challenging to take this into account in models that include a large number of receptor subunits. In our model, AMPAR subunits GluR1 and GluR2 can be in one of 21 or 5 states, respectively, when counting all the different phosphorylation states and bonds with other molecules (Tab. 1), which leads to 28^4^ possible types of tetramers. This makes it virtually impossible to model the dynamics of AMPAR tetramer assembly using the mass-action law-based approach where the concentration of each type of species is monitored (cf. [Michalski and Loew, 2012]). To avoid this problem, we used a statistical model that estimated the numbers and types of different types of AMPAR tetramers given the numbers of GluR1 and GluR2 subunits located at the membrane.

We assumed that the composition of AMPAR tetramers is random such that there is no preference of one type of subunit being more likely to bind with any other type of subunit. Thus, the probability of a tetramer being a GluR1 homomer without any S831-phosphorylated subunits is approximately

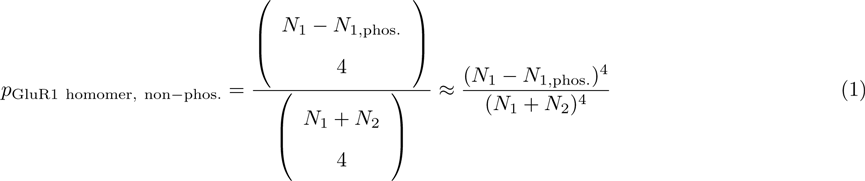

where *N*_1_ and *N*_2_ are the numbers of GluR1 and GluR2 subunits bound to the membrane, respectively, and *N*_1_,_phos_. is the number of S831-phosphorylated GluR1 subunits at the membrane (note that the *N*_1_,_phos_. subunits are included in all GluR1 subunits, i.e., *N*_1_,_phos_. ≤ *N*_1_). Accordingly, the probabilities of a tetramer being a GluR1 homomer with at least one S831-phosphorylated subunit, a GluR2 homomer, or a heteromer, are approximately:

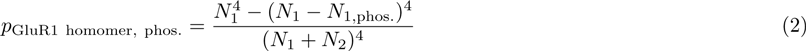

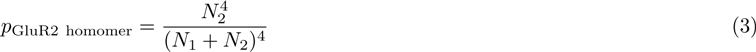

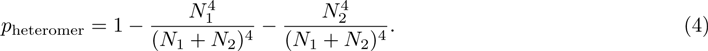

The number of tetramers that the *N*_1_ GluR1 subunits and *N*_2_ GluR2 subunits at the membrane can form is 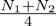. Here, we ignore the unpaired subunits by estimating that 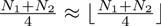. This gives us approximate values for expected numbers of different types of tetramers on the membrane:

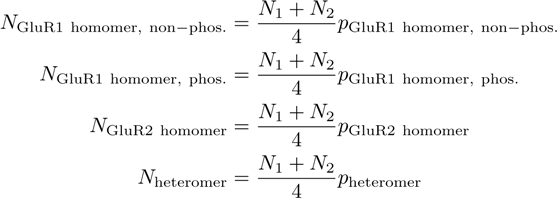

These estimates allow us to determine the total maximal synaptic conductance as the sum of the numbers of these tetramers multiplied with the corresponding single-channel conductances:

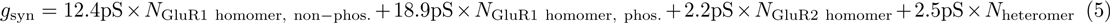

The single-channel conductance values 12.4 pS, 18.9 pS, 2.2 pS, and 2.5 pS are taken from experimental data [Oh and Derkach, 2005].

### 2.3 Modelling the Ca*^2+^* inputs and neuromodulatory inputs

We modelled the neurotransmission to the post-synaptic spine as fluxes of Ca^2+^ ions, *β*-adrenergic ligand, glutamate, and acetylcholine (labelled as Ca, L, Glu, and ACh, respectively, in Tab. 1). We used various stimulation paradigms: In Sections 3.2 and 3.5, long-lasting, single pulses of input species were applied. In Sections 3.3 and 3.6, we used the following repeated stimulus protocols: HFS — 100 pulses of Ca^2+^ (3 ms), repeated at 100 Hz; 4xHFS — 4 trains of HFS, separated by 3 seconds of quiescence; LFS — 900 pulses of Ca^2+^ (3 ms), repeated at 5 Hz. In these protocols, each Ca^2+^ pulse was accompanied by a 3-ms pulse of *β*-adrenergic ligand, glutamate, and acetylcholine, unless otherwise stated. In Section 3.7, the same approach was used, but the frequencies and numbers of repetitions of the inputs were taken from the experiments (see Tab. 4). Finally, in Section 3.4, we used a multicompartmental model of a layer 2/3 pyramidal cell [Markram et al., 2015] (L23 PC cADpyr229 5) to determine the amplitudes and time courses of the Ca^2+^ inputs conducted by NMDA receptors (NMDARs) when different pairing intervals were applied. We used the probabilistic AMPA–NMDA synapse model of [Hay and Segev, 2015] with the NMDA gating mechanism of [Spruston et al., 1995], a correction in the pre-synaptic resource update [Mäki-Marttunen et al., 2018] and an AMPA–NMDA ratio of 1:7. We estimated the numbers of Ca^2+^ ions entering into post-synaptic spines consisting of a 0.5 µm long and 0.1 µm thick neck and a 0.4 µm long and 0.4 µm thick head across time, and used these numbers as the input to the biochemical model. The effects of LTP/LTD on the size of Ca^2+^ inputs were not considered in this work.

### 2.4 Parameter alterations and model fitting

In Sections 3.5 and 3.7, we altered the initial concentrations of many proteins to explore the parameter space or to perform model fitting. The concentrations of upstream PKA-pathway proteins R (*β*-adrenergic receptor), Gs, AC1, and AC8 were varied in proportion using a coefficient *f*_PKA_ *∈* [0, 2], and, likewise, the concentrations of upstream PKC-pathway proteins mGluR, M1, Gq, and PLC using a coefficient *f*_PKC_ *∈* [0, 2]. Furthermore, in Section 3.7, calmodulin and CaMKII were altered in proportion by a factor *f*_CaMKII_ *∈* [0, 2], phosphatases PP1 and PP2B by a factor *f*_PP_ *∈* [0, 2], and phosphodiesterases PDE1 and PDE4 by a factor *f*_PDE_ *∈* [0, 2]. In both sections, the rapidity of Ca^2+^ expulsion was varied by altering the concentration of NCX, and in Section 3.7, the concentrations of PKA and PKC were varied in addition to the upstream proteins — these concentrations were varied within the interval from 0 to double the original value. For the multi-objective optimisation in Section 3.7, we used the Python implementation (published by the authors of [Bahl et al., 2012]) of the non-dominated sorting genetic algorithm II (NSGA-II) [Deb et al., 2002] with population size 1000. To restrict to physiologically realistic Ca^2+^ dynamics, we disregarded the data where free Ca^2+^ concentrations rose above 2 µM for one or more levels of Ca^2+^ input in Section 3.5. In a similar manner, in Section 3.7, we introduced an objective function that penalised parameter sets that produced Ca^2+^ transients larger than 2 µM.

### 2.5 Simulation software and code accessibility

For deterministic simulations of intracellular signalling, we used the NEURON simulator with the reaction-diffusion (RxD) extension [McDougal et al., 2013]. For stochastic simulations, we used NeuroRD software (https://github.com/neurord). In both types of simulations, we used adaptive time-step integration methods. The NEURON simulator was also used for simulating the multicompartmental model of layer 2/3 pyramidal cell in Section 3.4. The full model along with the fitting and data-analysis algorithms (Python scripts) that were used in this study are publicly available in ModelDB at http://modeldb.yale.edu/260971 (password “plasticity” required during peer review).

## 3 Results

### 3.1 Literature review and model construction

We reviewed the literature of long-term plasticity in the neocortex, focusing on the molecular signalling pathways that are needed for LTP/LTD in the post-synaptic spine of pyramidal cells (Tab. 3A). Three main pathways were highlighted in the experimental studies, namely, the PKA, PKC, and CaMKII pathways. To construct a computational model of cortical post-synaptic plasticity that describes these pathways, we adopted mass-action law-based descriptions of these pathways from biochemically detailed models of post-synaptic LTP/LTD in other brain areas, namely, hippocampus, basal ganglia and cerebellum (Tab. 3B). We prioritised the model components from hippocampal models due to the relatively small ontological differences between hippocampus and neocortex [Kirsch and Chechik, 2016]. We focused on the effects of these pathways on AMPARs due to the better description of intracellular regulation of AMPAR dynamics in comparison to that of NMDA and kainate receptors or voltage-gated ion channels. In short, we based our model on that of [Jedrzejewska-Szmek et al., 2017], which describes the PKA- and CaMKII-dependent forms of GluR1 phosphorylation, and added the mGluR- and M1-mediated activation of PKC from [Kim et al., 2013] and [Blackwell et al., 2018], respectively. We then adopted the reactions describing PKC-dependent endocytosis of GluR2 and reinsertion to the membrane from [Gallimore et al., 2018], which allowed the representation of post-synaptic depression with our model. The pathways included in the model are illustrated in Fig. 2. A description of the model calibration is given in Section 2.1, and the full set of model reactions and initial concentrations is provided in Tables 1 and 2, respectively.

**Figure 2:**
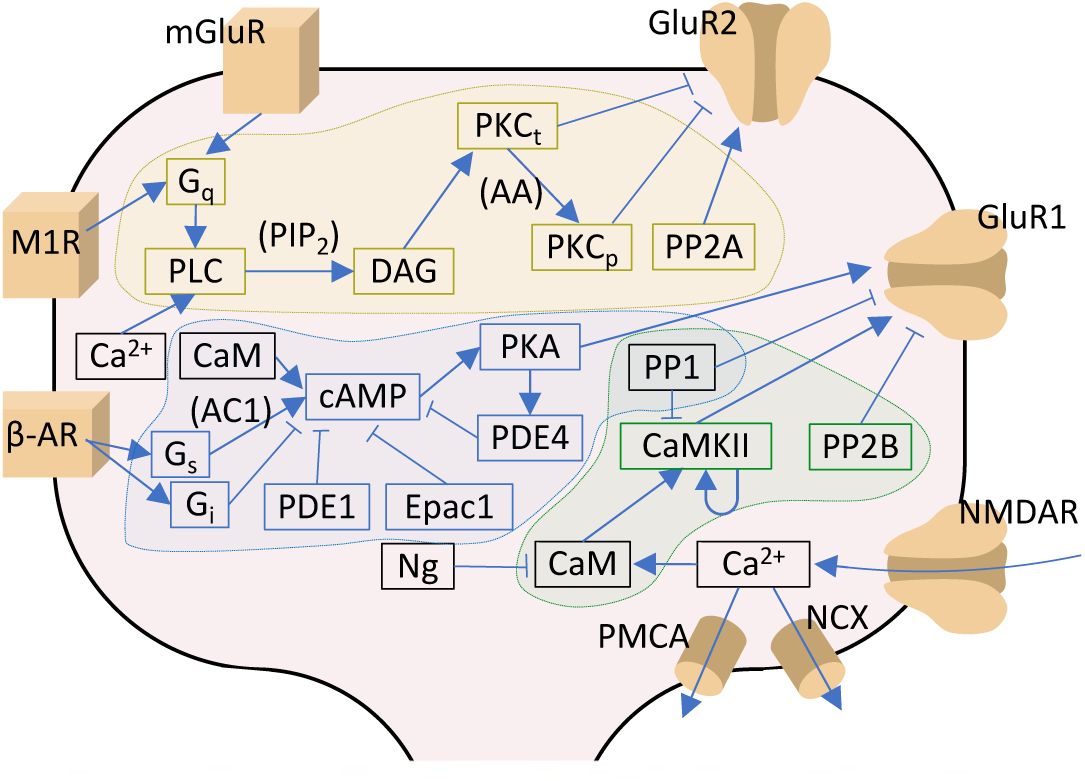
Signalling pathways included in the model. The PKA-pathway-related proteins and signalling molecules are highlighted by blue, PKC-pathway molecules by yellow, and CaMKII-pathway molecules by green colours. Reactions associated with a molecular species in parenthesis indicate a dependency on the denoted species — for details, see Tab. 1. *β*-AR – *β*-adrenergic receptor; AC1 & AC8 – adenylyl cyclase type 1 or 8; CaM – calmodulin; CaMKII – calmodulin-dependent protein kinase II; cAMP – cyclic adenosine monophosphate; DAG – diacylglycerol; Epac1 – exchange factor directly activated by cAMP 1; Gi, Gq & Gs – G-protein type I, Q, or S; GluR1 & GluR2 – AMPAR subunit 1 or 2; mGluR – metabotropic glutamate receptor; M1R – cholinergic receptor M1; NCX – Na^+^-Ca^2+^ exchanger; Ng – neurogranin; NMDAR – NMDA receptor; PDE1 & PDE4 – phosphodiesterase type 1 or 4; PIP_2_ – phosphatidylinositol 4;5-bisphosphate; PKA – protein kinase A; PKCt & PKCp – transiently or persistently active protein kinase C; PLC – phospholipase C; PMCA – plasma membrane Ca^2+^ ATPase; PP1 – protein phosphatase 1; PP2A – protein phosphatase 2A; PP2B – protein phosphatase 2B (calcineurin). In this work, the NMDARs are considered only in Section 3.4: in the rest of the work, Ca^2+^ is directly injected as a square-pulse current into the spine.

**Table 3:** Pathways contributing to cortical synaptic plasticity. **A**: Experimental evidence on the requirement of various molecular species for specific types of synaptic regulation in different cortical areas. **B**: Model components needed for describing the modes of plasticity listed in (A). References are made to previous computational models describing these pathways. The types of phosphorylation of AMPAR subunit that mediate the plasticity are printed in bold.

| **A** |  |  |  |  |
| --- | --- | --- | --- | --- |
| Pathway components | Type of neurons | Type of regulation | Pre-/post-syn. | References |
| CaMKII | Cingulate cortex | Esophageal acid-induced sensitisation | post-syn. | [Banerjee et al., 2013] |
| CaMKII | Prefrontal cortex, pyramidal neurons | 5-HT1-induced modulation of AMPA currents | post-syn. | [Cai et al., 2002] |
| β-adr. receptors, PKA | Visual cortex, layer 4 pyramidal cells | Potentiation of AMPA currents | post-syn. | [Seol et al., 2007] |
| M1 receptors, PKC | Visual cortex, layer 4 pyramidal cells | Depression of AMPA currents | post-syn. | [Seol et al., 2007] |
| D1-PKA | Prefrontal cortex, pyramidal neurons | Potentiation of AMPA currents | post-syn. | [Sun et al., 2005] |
| β-adr. receptors | Frontal cortex | Potentiation of field EPSPs | n/a | [Sáez-Briones et al., 2015] |
| PKC | Cultured cortical neurons | Internalisation of AMPARs | post-syn. | [Chung et al., 2000] |
| ERK | Visual cortex | Potentiation of field EPSPs | n/a | [Di Cristo et al., 2001] |

| **B** |  |
| --- | --- |
| Molecular pathway | Cell type and references |
| $\text{Ca}^{2+} \rightarrow \text{CaM} \rightarrow \text{CaMKII}$ | Hippocampal CA1 neuron [Bhalla and Iyengar, 1999, Jedrzejewska-Szmek et al., 2017], generic [Hayer and Bhalla, 2005], cerebellar Purkinje cells [Gallimore et al., 2018], striatal spiny projection neuron [Blackwell et al., 2018] |
| $\text{CaMKII} \rightarrow \text{GluR1 S831p}$ | Hippocampal CA1 neuron [Jedrzejewska-Szmek et al., 2017] |
| β-adrenergic receptors → cAMP<br>cAMP → PKA | Hippocampal CA1 neuron [Jedrzejewska-Szmek et al., 2017]<br>Hippocampal CA1 neuron [Bhalla and Iyengar, 1999, Jedrzejewska-Szmek et al., 2017], cerebellar Purkinje cells [Gallimore et al., 2018] |
| $\text{PKA} \rightarrow \text{GluR1 S845p}$ | Hippocampal CA1 neuron [Jedrzejewska-Szmek et al., 2017] |
| M1 receptors → PLC<br>PLC → PKC | Cerebellar Purkinje cells [Gallimore et al., 2018]<br>Hippocampal CA1 neuron [Bhalla and Iyengar, 1999], striatal spiny projection neuron [Kim et al., 2013, Blackwell et al., 2018]<br>cerebellar Purkinje cells [Hellgren Kotaleski et al., 2002, Gallimore et al., 2018] |
| $\text{PKC} \rightarrow \text{GluR2 S880p}$ | Cerebellar Purkinje cells [Gallimore et al., 2018] |

**Table 4:**
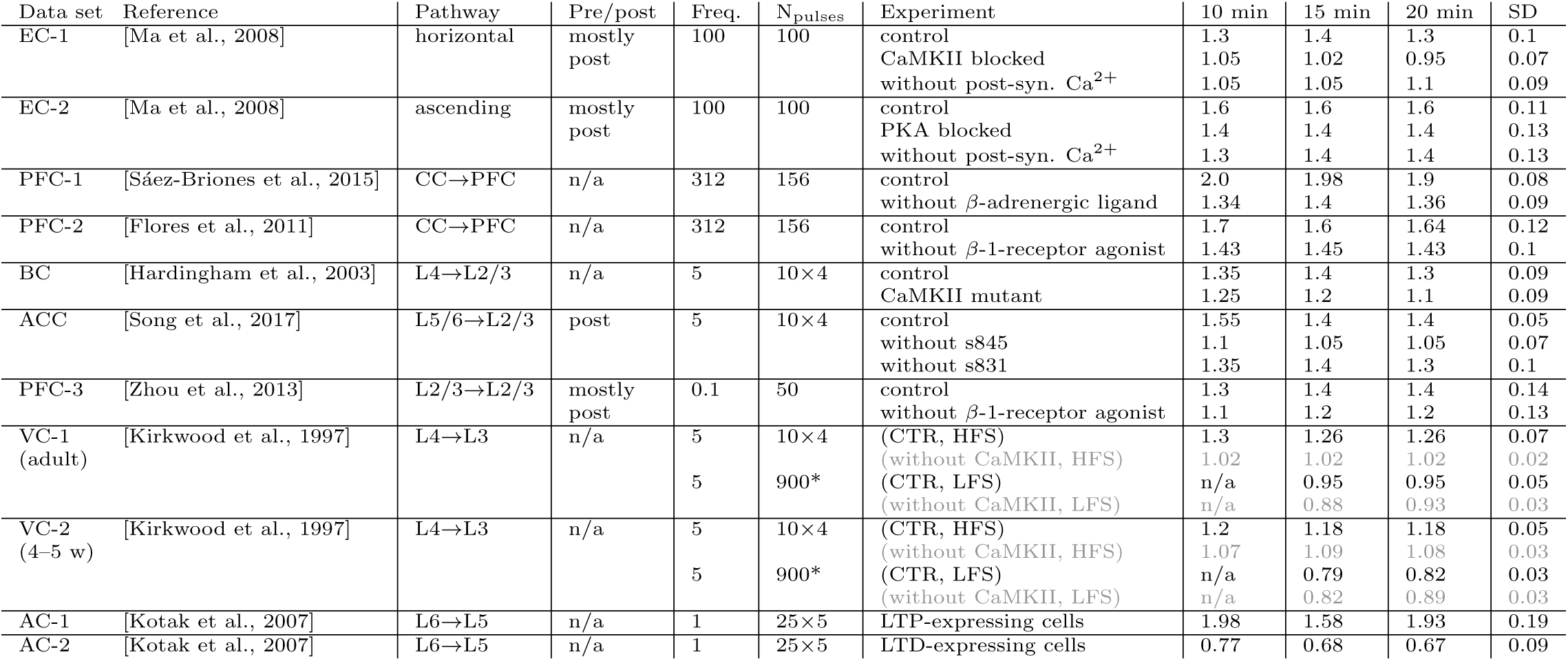
List of LTP/LTD experiments in the cortex. The first column labels the experimental data set and names the underlying study. The second column shows the considered synaptic pathway and the third column shows whether the observed LTP/LTD had a pre- or post-synaptic origin. The fourth and fifth columns show the frequency (in Hz) of stimulation and the number of pulses delivered, respectively: 10*×*4 means that 10 trains of 4 pulses with 10 ms interval (100 Hz) were delivered, and likewise, 25*×*5 means that 25 trains of 5 pulses with 10 ms interval were delivered. The sixth column tells whether the data were obtained in control conditions or under additional blockers or agonists. The seventh, eighth, ninth, and tenth columns show the relative change in synaptic strength 10, 15, and 20 min after the start of the stimulus protocol and an average SD of the relative synaptic strengths — these values were approximated from the LTP/LTD curves plotted in the underlying references. The rows correspond to experiments from a given reference that are divided to 11 different experimental data sets. Within each data set, the underlying system is assumed to be otherwise similar to the control except for the applied modifier: as an example, the chemical or genetic blockade of CaMKII activity (as performed in [Ma et al., 2008] and [Hardingham et al., 2003]) is here expected to only affect the ability of CaMKII to autophosphorylate, and the rest of the model parameters are kept fixed. The experiments printed in grey were included in the underlying study, but were excluded from the main analyses of the present work (see main text). EC – entorhinal cortex; PFC – prefrontal cortex; BC – barrel cortex; ACC – anterior cingulate cortex; VC – visual cortex; AC – auditory cortex; CC – corpus callosum. (*): The LFS of 900 3-ms pulses at 5 Hz in data sets VC-1 and VC-2 was replaced by 180 15-ms pulses at 1 Hz to decrease computational load in the optimisation.

### 3.2 Ca*^2+^* activates multiple pathways that regulate the post-synaptic plasticity in cortical PCs

All pathways of Tab. 3B are Ca^2+^-dependent, but due to the variability in binding rates and quantities of different Ca^2+^-binding molecules, some pathways become more easily activated than others. This permits LTP or LTD to be induced in a way that is sensitive to the amount of Ca^2+^ inputs and may serve as a basis for BCM-type rules of plasticity.

To examine the sensitivities of LTP- and LTD-inducing pathways to Ca^2+^, we simulated the injection of a prolonged square-pulse Ca^2+^ input of varying magnitude (illustrated in Fig. 3A) into the post-synaptic spine and quantified the degree of activation of each of the Ca^2+^-binding molecules and the downstream signalling cascades. The simulations were carried out in the presence of mGluRs and *β*-adrenergic and cholinergic neuromodulation, which were modelled as prolonged square-pulse inputs as well.

**Figure 3:**
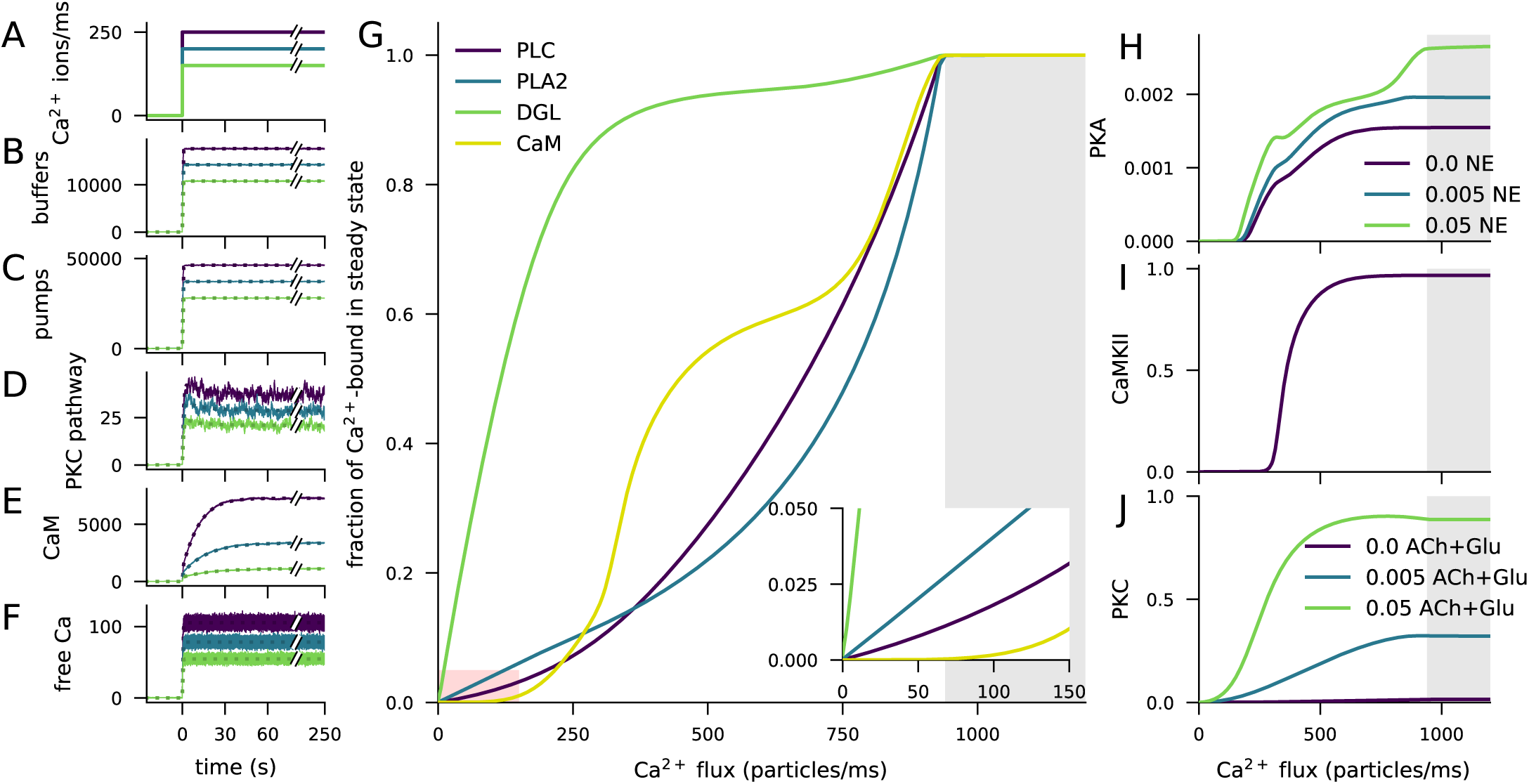
Ca*^2+^* activates CaMKII, PKA, and PKC pathways. **A**: Illustration of the stimulus protocols with Ca^2+^ flux amplitudes 150 (green), 200 (cyan), and 250 (purple) particles/ms. **B–F**: Time courses of Ca^2+^ (in #particles) bound to buffers (B), pumps (C), PKC-pathway proteins (D), or CaM (E), and the concentration of free Ca^2+^ ions (F), according to NeuroRD (solid; averaged across 8 samples) or NEURON (dashed) simulations. Colours indicate the Ca^2+^ flux used (see (A)). **B**: Number of Ca^2+^ ions bound to Ca^2+^ buffers, i.e., immobile buffer and calbindin. **C**: Number of Ca^2+^ ions bound to Ca^2+^ pumps and exchangers, i.e., PMCA and NCX. **D**: Number of Ca^2+^ ions bound to PKC-pathway proteins PLC and PLA2. **E**: Number of Ca^2+^ ions bound to CaM, in all its forms. **F**: Cytosolic Ca^2+^ concentration (mM) **G**: Degrees of activation of different Ca^2+^-binding proteins in a steady state (5 min after onset of Ca^2+^ input) as a function of the magnitude of Ca^2+^ flux. The x-axis shows the amplitude of the Ca^2+^ input (see panel (A)), and the y-axis shows the ratio of the underlying species in a Ca^2+^-bound form over the total number of the proteins. For CaM, only the CaM molecules bound by four Ca^2+^ ions are considered activated — in PLC, PLA2, and DGL, binding of only one Ca^2+^ ion is needed for activation. Here, the measured quantity of active PLC includes both Gq-bound and non-Gq-bound CaPLC. Inset: zoomed-in view on the red area. **H**: Ratio of the steady-state concentration of PKA catalytic subunit over the theoretical maximum where all PKA molecules were dissociated into residuals and catalytic subunits. Colour of the curve indicates the amplitude of the *β*-adrenergic ligand flux (particles/ms). **I**: Fraction of phosphorylated CaMKII subunits. **J**: Fraction of (transiently or persistently) activated PKC. Colour of the curve indicates the amplitude of the cholinergic and glutamatergic ligand flux (particles/ms). The grey area in panels (G)–(J) represents Ca^2+^ inputs that cause cytosolic Ca^2+^ concentration to reach extremely high levels (*>* 1 mM) that are likely to lead to apoptosis.

The injected Ca^2+^ quickly bound to Ca^2+^ buffers (immobile buffer and calbindin, Fig. 3B) and pumps (PMCA and NCX, Fig. 3C) as well as to the proteins of the PKC pathway (PLA2 and PLC, Fig. 3D): a 95-% saturation was reached in 1–2 s (Fig. 3B–D). In contrast, the activation of CaM was slower (Fig. 3E): a 95-% saturation was reached in 32–53 s, depending on the magnitude of the Ca^2+^ input. Consistent with experimental literature, a vast majority of Ca^2+^ was quickly bound and only a small fraction remained free in the cytosol (Fig. 3F).

To further illustrate the differences between the activation patterns of these pathways, we quantified the degrees of Ca^2+^ binding of these molecules in a steady state (5 min after the onset of Ca^2+^) and the overall activation/deactivation of downstream molecules as a function of the magnitude of the Ca^2+^ injection. Both PKC pathway-mediating proteins PLC, DGL, and PLA2, along with the PKA and CaMKII pathway-related protein CaM became completely activated if large enough Ca^2+^ flux is given, but their degrees of activity varied across the magnitude of the injected Ca^2+^ flux (Fig. 3G). DGL was most completely activated throughout the Ca^2+^ amplitude, owing to the large equilibrium constant of its Ca^2+^ binding. At extremely large Ca^2+^ fluxes, CaM was more completely bound by Ca^2+^ than PLC and PLA2 (Fig. 3G), but at lower Ca^2+^ amplitudes, CaM remained largely unbound (Fig. 3G inset). This is reflected in the activation patterns of the catalytic subunit of PKA (Fig. 3H) and CaMKII (Fig. 3I), both of which are dependent on the activation of CaM and thus had a small response at low Ca^2+^ amplitudes. PKC, by contrast, became activated at relatively small Ca^2+^ amplitudes (Fig. 3J). Of these three pathways, the PKC pathway was dependent on the cholinergic ligands or the activation of the mGluRs (Fig. 3J), and the PKA pathway was dependent on the availability of *β*-adrenergic ligands (Fig. 3H). Taken together, these results highlight the need for large Ca^2+^ flux to the post-synaptic spine for the activation of the CaMKII pathway, relatively large Ca^2+^ flux for the activation of the PKA pathway, and relatively small Ca^2+^ flux for the activation of the PKC pathway.

### 3.3 High-frequency stimulation causes LTP and low-frequency stimulation causes LTD in GluR1-GluR2-balanced synapses

The Ca^2+^ flux entering the post-synaptic spine is extremely large during and after synaptic transmission and low otherwise, which causes the signalling pathways to be activated and deactivated in a more dynamic way than described in the Section 3.2. The activation of these pathways and their dependence on the stimulus protocol are difficult to study experimentally due to methodological constraints (e.g., side effects of fluorescence indicators, lack of signal calibration, and poor temporal or spatial resolution), but biochemically detailed models, such as the one considered in this work, can provide insights into the transient molecular mechanisms behind LTP and LTD. Our model is particularly well suited to study the mechanisms behind CaMKII-, PKA- and PKC-mediated phosphorylation of AMPAR subunits, which are important mediators of long-term plasticity [Wang et al., 2005]. Phosphorylation of GluR1 subunits at S845 increases the insertion rate of the AMPAR into the membrane, thus leading to post-synaptic LTP [Diering et al., 2016]. Conversely, phosphorylation of GluR2 subunits at S880 increases the rate of receptor endocytosis from the membrane, and has thus been observed to lead to post-synaptic LTD [Xia et al., 2000]. However, it is not the number of the membrane-expressed AMPAR subunits alone that determine the strength of the synapse, but different compositions of the subunits have different single-channel conductances, and phosphorylation at S831 of the GluR1 subunit also affects the conductance of the channel [Oh and Derkach, 2005].

To simulate the reaction dynamics in the post-synaptic spine under realistic input patterns, we applied the 4xHFS and LFS protocols. Each input contained transient (3 ms) influxes of Ca^2+^ (1900 particles/ms) into the cytosol and glutamate (20 particles/ms), acetylcholine (20 particles/ms) and *β*-adrenergic ligand (10 particles/ms) into the extracellular subspace near the spine membrane. We used a balanced ratio (1:1) of GluR1 and GluR2 subunits. We recorded the time courses of the concentrations of all CaMKII-, PKA-, and PKC-pathway molecules contributing to LTP or LTD to monitor their activity during and following the stimulation protocols. We also recorded the numbers of membrane-inserted GluR1 and GluR2 and their state of phosphorylation and used Eq. 5 for determining the maximal synaptic conductance.

In the 4xHFS protocol, which typically causes LTP in plasticity experiments, our model predicts a large increase in total synaptic conductance (Fig. 4A) due to a radical increase in membrane-inserted GluR1 subunits (Fig. 4B) and a decrease in GluR2 subunits (Fig. 4C). These changes in membrane-expression of AMPAR subunits were dependent on activations of many signalling proteins in the CaMKII (Fig. 4D–H), PKA (Fig. 4I–M), and PKC (Fig. 4N–R) pathways. First, the Ca^2+^ entry (Fig. 4D) caused a rapid increase in half-activated calmodulin (bound by two Ca^2+^ ions; Fig. 4E), leading to a longer-lasting increase in active calmodulin (Fig. 4F). Calmodulin activation led to an increase in the concentration of phosphorylated CaMKII (Fig. 4G), which phosphorylated the GluR1-type receptors at S831 (Fig. 4H). The *β*-adrenergic input (Fig. 4I), in turn, bound to the *β*-adrenergic receptors and activated the Gs proteins (Fig. 4J), which bound to the adenylyl cyclase AC1 to produce cAMP (Fig. 4K). cAMP bound to PKA to release the catalytic subunits of PKA (Fig. 4L), which led to a phosphorylation of the GluR1-type receptors at S845 (Fig. 4M) and thus to increased membrane expression of GluR1 subunits and total synaptic conductance (Fig. 4A–B). Due to the simultaneous activation of the CaMKII pathway, a significant proportion of double phosphorylated GluR1-type receptors was observed (Fig. 4H, M). As for the PLC–PKC pathway, glutamate (Fig. 4N, blue) bound to mGluRs and acetylcholine (Fig. 4N, green) bound to muscarinic receptors (M1), and the activation of these receptors contributed to the activation of Gq proteins (Fig. 4O). The activated Gq proteins bound with phospholipase C (PLC) and metabolised phosphatidylinositol 4,5-bisphosphate (Pip2) into diacylglycerol (DAG, Fig. 4P), which activated protein kinase C (PKC, Fig. 4Q). This led to the phosphorylation of GluR2-type receptors at S880 (PKC, Fig. 4R), which caused the decrease in membrane expression of GluR2 observed in Fig. 4C.

**Figure 4:**
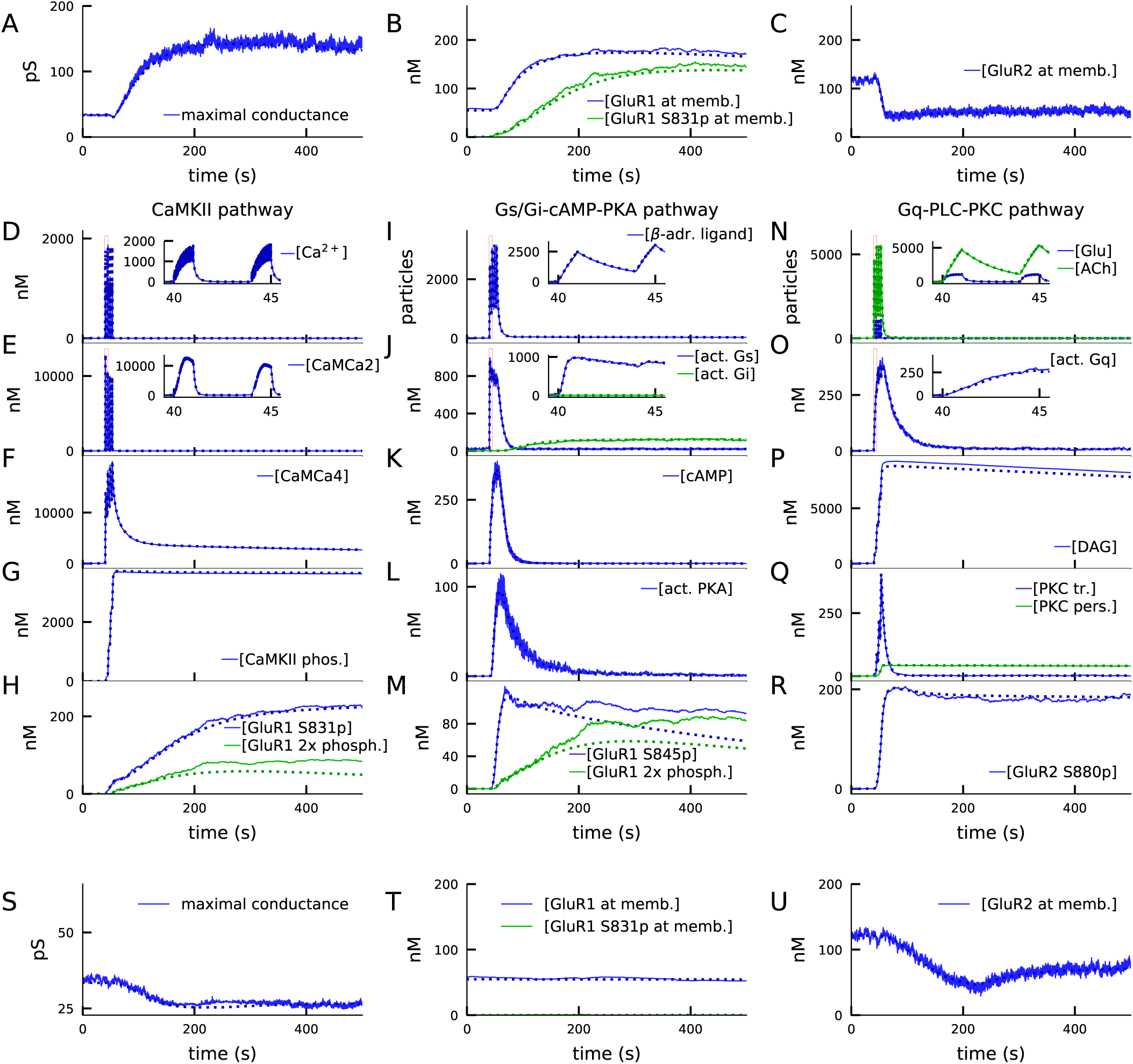
4xHFS activates CaMKII, PKA, and PKC pathways and leads to LTP (A–R), while LFS activates the PKC pathway and leads to LTD (S–U). **A**: Total synaptic conductance in response to 4xHFS, determined by the numbers of membrane-inserted GluR1s and GluR2s — see Eq. 5. The stimulation starts at 40 s and lasts until 53 s. **B–C**: Concentration of membrane-inserted GluR1s (B) and GluR2s (C) in response to 4xHFS. **D–H**: Concentration of different species in the CaMKII pathway, namely, intracellular unbound Ca^2+^ (D), CaM bound with two Ca^2+^ ions (E), CaM bound with four Ca^2+^ ions (active CaM; F), phosphorylated CaMKII, bound or unbound by CaMCa4 (G), and S831-phosphorylated and double-phosphorylated GluR1 subunits (H) in response to 4xHFS. **I–M**: Concentration of different species in the cAMP-PKA pathway, namely, *β*-adrenergic ligand in all its forms (I), activated (GTP-bound but not bound to ATP) Gs and Gi proteins (J), intracellular cAMP (K), catalytic subunit of PKA (L), and S845-phosphorylated and double-phosphorylated GluR1 subunits (M) in response to 4xHFS. **N–R**: Concentration of different species in the PLC-PKC pathway, namely, glutamate and acetylcholine in all their forms (N), activated (GTP-bound but not bound to DAG) Gq proteins (O), intracellular DAG (P), activated PKC (Q), and S880-phosphorylated GluR2 subunits (R) in response to 4xHFS. **S**: Total synaptic conductance in response to LFS. **T–U**: Concentration of membrane-inserted GluR1s (T) and GluR2s (U) in response to LFS, which starts at 40 s and lasts until 220 s. The solid lines represent stochastic (NeuroRD) simulation results, while the dashed lines represent data from deterministic (NEURON RxD) simulations. *β*-adrenergic ligands, glutamate, and acetylcholine are measured in numbers of particles as they reside both at the membrane (when bound to receptors) and at the extracellular subspace near the spine membrane (when unbound); other species measured in concentration.

The differences between NEURON and NeuroRD simulation results in Fig. 4M were due to the stochasticity in NeuroRD simulator — both smaller and larger GluR1 phosphorylation levels compared to NEURON simulation results (Fig. 4M, dashed) were obtained when NeuroRD simulations were run with different random number seeds (not shown).

In the LFS protocol, which typically causes LTD in the experiments, our model predicts a prominent (20%) decrease in total synaptic conductance (Fig. 4S) due to a decrease in GluR2 subunits. In this protocol, the Ca^2+^ inputs are insufficiently large to activate CaM, and the Gs proteins remain deactivated as well (data not shown). In consequence, CaMKII and PKA pathways remain deactivated, and the effect of the LFS protocol on GluR1 phosphorylation and membrane insertion is small (Fig. 4T). By contrast, the PKC pathway is almost as strongly activated as in the 4xHFS protocol (data not shown), leading to prominent S880 phosphorylation of GluR2 (data not shown) and removal of GluR2 from the membrane (Fig. 4U).

The expression of both LFS-induced LTD and 4xHFS-induced LTP of these types is dependent on the presence of both GluR1 and GluR2 subunits: GluR1-deficient synapses failed to show 4xHFS-induced LTP (extended data set, Fig. 4.1A) and GluR2-deficient synapses failed to show LFS-induced LTD (Fig. 4.1B). To show that our results were not an artefact of the tetramer formation rule (Eq. 1–5), we applied an alternative tetramer formation rule where GluR1 and GluR2 subunits randomly dimerised and the dimers paired with like dimers (which disallows the emergence of heterotetramers with 1:3 or 3:1 proportion of GluR1:GluR2 subunits; cf. [Gan et al., 2015]). We reproduced the LFS-induced LTD and 4xHFS-induced LTP using this dimer-of-like-dimers rule with a modified (35:65) balance of GluR1 vs. GluR2 subunits (Fig. 4.2A).

The activations of the above pathways are dependent on the magnitude and dynamics of the inputs to the model, namely, Ca^2+^, *β*-adrenergic and cholinergic ligands, and glutamate. All pathways leading to GluR1 and GluR2 phosphorylation and the consequent exocytosis and endocytosis are Ca^2+^-dependent: blocking Ca^2+^ entry completely abolished 4xHFS-induced LTP (Fig. 5A) that followed GluR1 insertion (Fig. 5B) and GluR2 endocytosis (Fig. 5C). Blocking *β*-adrenergic ligands abolished the 4xHFS-induced LTP (Fig. 5A) by suppressing the membrane-insertion of GluR1 (Fig. 5B), but had no effect on GluR2 endocytosis (Fig. 5C). Likewise, blocking *β*-adrenergic ligands had no effect on LFS-induced LTD (not shown). In contrast, LFS-induced LTD (Fig. 5E) that followed GluR2 endocytosis (Fig. 5G) was reduced by blockade of mGluR activation while the number of GluR1 subunits at the membrane remained unaffected (Fig. 5F). This reduction was strengthened by simultaneous blockade of cholinergic inputs (Fig. 5E–G, yellow traces). Counterintuitively, blocking mGluR and M1-receptor activation also reduced the amplitude of the 4xHFS-induced LTP (Fig. 5A) by disabling GluR2 endocytosis (Fig. 5C) while it had no effect on GluR1 insertion (Fig. 5B). The reason for this is that in the PKC pathway-blocked case there is a smaller post-4xHFS membrane-bound GluR1 ratio (fraction of GluR1 subunits over all GluR subunits at the membrane) than in the control case, and thus the probability of AMPARs being homomeric GluR1 tetramers (which had a very large conductance compared to other tetramers; Eq. 5) is much smaller in the former case than in control (Fig. 5D). Although qualitatively similar difference can be observed in post-LFS membrane-bound GluR1 ratios between PKC pathway-blocked case and control, the probability of homomeric GluR1 tetramers is very small in both cases (Fig. 5H) and thus the LFS-induced LTD is not affected.

**Figure 5:**
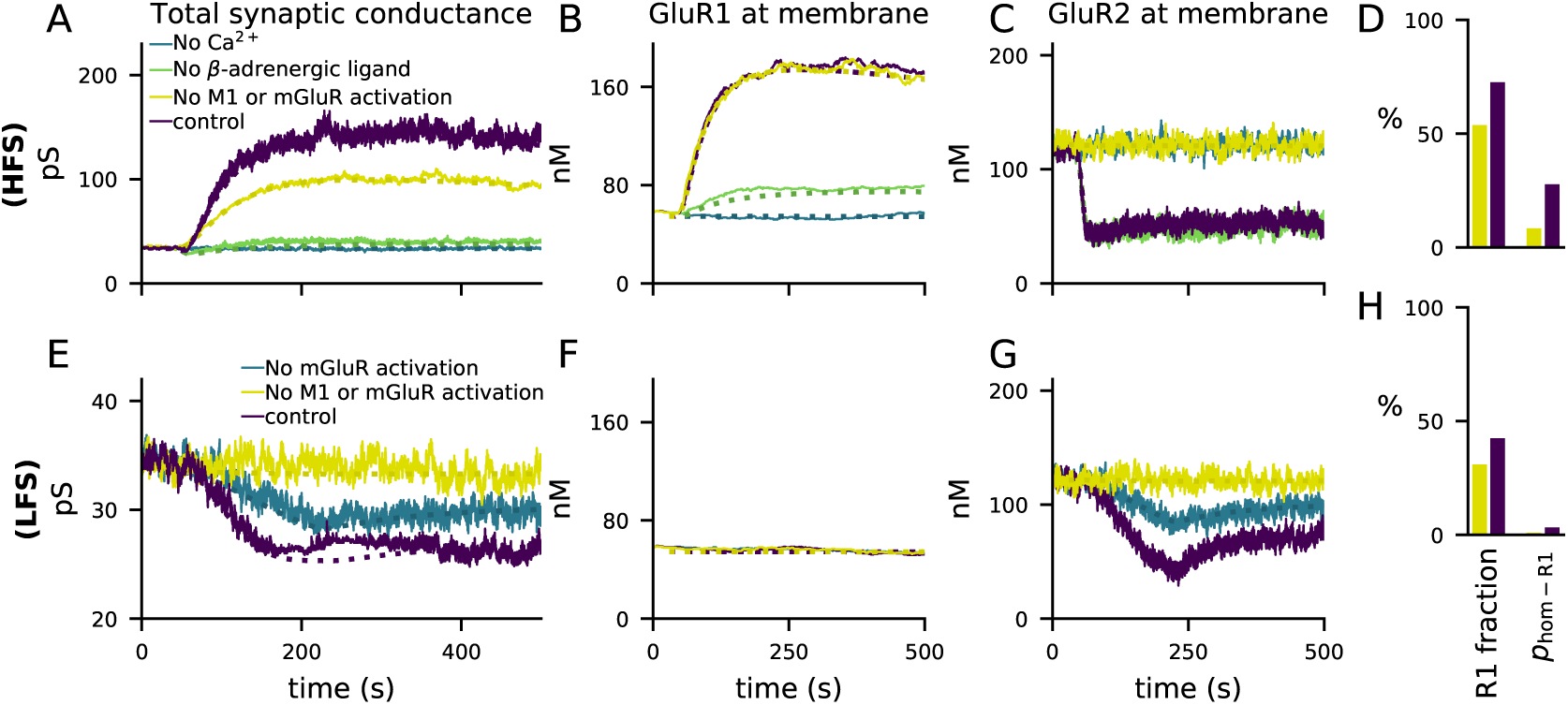
4xHFS-induced LTP is dependent on *β*-adrenergic ligands and LFS-induced LTD is dependent on activation of mGluRs or cholinergic receptors. **A–D**: 4xHFS-induced LTP in the control case (dark purple), without Ca^2+^ inputs (blue), without *β*-adrenergic ligands (green), and under blockade of PKC pathway-activation (mGluRs or cholinergic receptors; yellow). **E–H**: LFS-induced LTD in the control case (dark purple), under the blockade of mGluR activation (blue), and under blockade of both mGluRs or cholinergic receptors (yellow). **A, E**: Total synaptic conductance. **B, F**: Membrane expression of GluR1. **C, G**: Membrane expression of GluR2. **D, H**: The fraction of membrane-inserted GluR1 over all membrane-inserted GluR subunits (left) and the probability of an AMPAR tetramer being homomeric GluR1 (right) at the end of the 4xHFS (D) or LFS (H) simulation with (dark purple) and without (yellow) PLC-activating ligands.

Taken together, our results show that cortical synapses expressing both GluR1 and GluR2 subunits can express a frequency-dependent form of post-synaptic plasticity (LTP for high-frequency inputs, LTD for low-frequency inputs) that is gated by neuromodulators affecting the PKA and PKC pathways. Our findings also lend support to that GluR2 endocytosis may lead to either potentiation (Fig. 5A) or depression (Fig. 5E), depending on the prevalence of the GluR1 subunits at the membrane.

### 3.4 Paired-pulse stimulus protocol induces PKA- and PKC-dependent spike-timing-dependent plasticity (STDP) in GluR1-GluR2-balanced synapses

Cortical synapses typically exhibit a type of synaptic plasticity, namely STDP, that is dependent on both the pre- and post-synaptic activity. According to a classical model, the differences in the outcome of STDP for different pairing intervals of pre- and post-synaptic stimulus are explained by different amount of Ca^2+^ entering the post-synaptic spine, which is affected by both the pre-synaptically released glutamate and the elevation of post-synaptic membrane potential. Biophysically detailed neuron modelling offers a powerful tool for determining the size of these Ca^2+^ inputs as a function of the pairing interval.

We considered the LTP/LTD response to paired stimulation protocol using a multicompartmental model of a layer 2/3 pyramidal cell (Fig. 6A) [Markram et al., 2015]. We placed a synaptic spine with volume 0.5 µm^3^ at a random location on the apical dendrite, 250-300 µm from the soma (Fig. 6A, thick, black branches), and stimulated the head of the spine with glutamatergic synaptic currents [Hay and Segev, 2015, Markram et al., 2015] (Fig. 6A, black traces, top). In parallel, we stimulated the soma with a burst of four short (2 ms) supra-threshold square-pulse currents (Fig. 6A, black traces, bottom). Given that approximately 10% of the NMDAR-mediated currents and none of the AMPAR-mediated currents are conducted by Ca^2+^ flux, we could determine the number of Ca^2+^ ions entering the spine at each time instant following the onset of the synaptic input (Fig. 6A, grey traces). This experiment was repeated using different inter-stimulus intervals (ISI) between the synaptic and somatic stimuli and averaged across *N_samp_* = 30 trials, yielding a unique Ca^2+^ flux time course for each pairing ISI (Fig. 6B–D). These Ca^2+^ flux time series were imported into our biochemical model, which allowed us to predict the magnitude of GluR subunit phosphorylation and membrane insertion for each pairing interval. Throughout these experiments, the activation of mGluRs was blocked.

**Figure 6:**
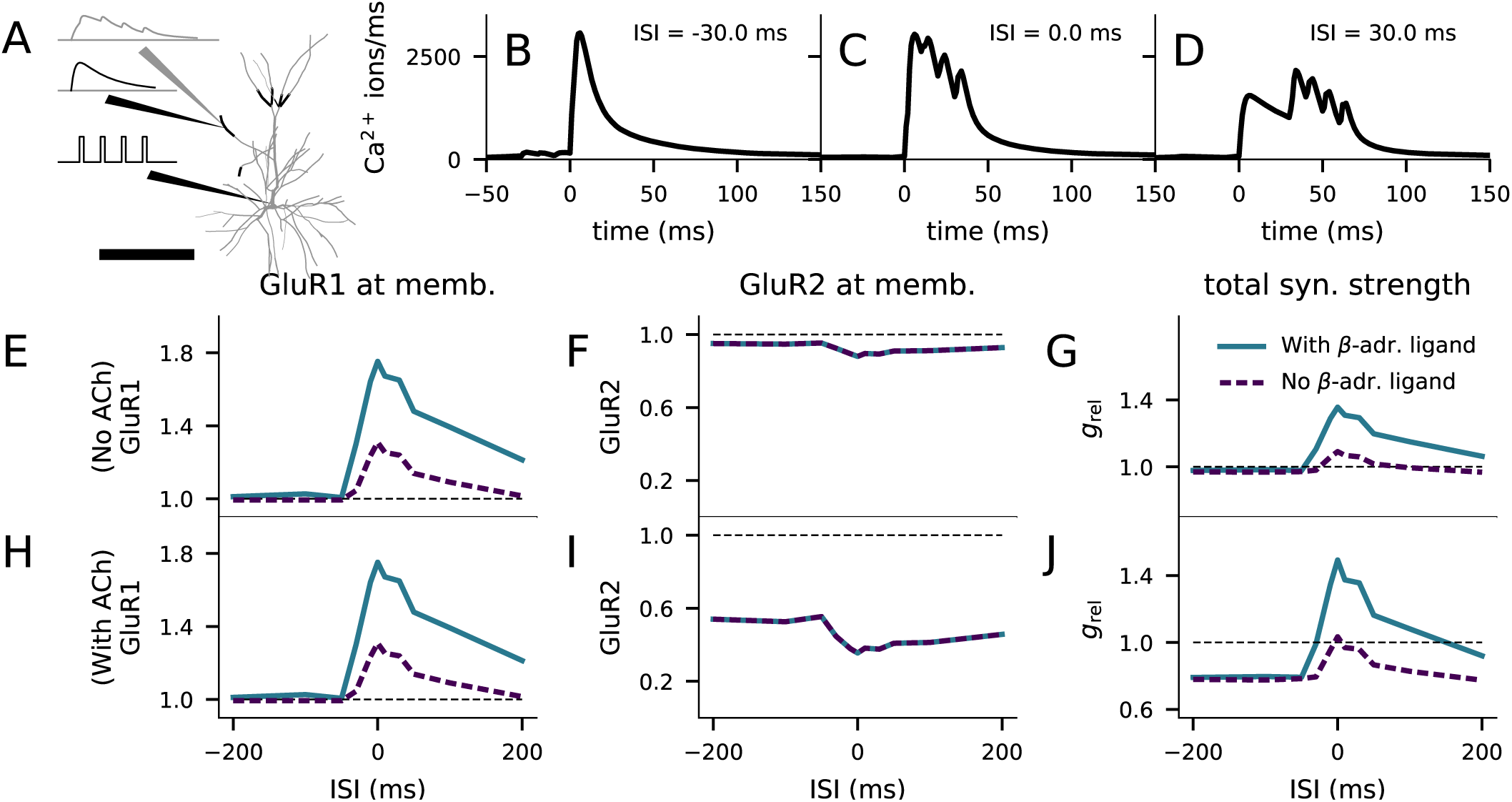
Layer 2/3 pyramidal cell plasticity in response to STDP protocol depends on neuromodulatory state and pairing interval. **A**: Layer 2/3 pyramidal cell morphology (grey, thin), locations of synaptic input highlighted (black, thick). Inset: Illustration of the inputs (black) and the recorded synaptic intracellular Ca^2+^ (grey). Scale bar 200 µm. **B–D**: Ca^2+^ flux to the dendritic spine when the somatic stimulation onset is 30 ms prior to (B), at the same time as (C), or 30 ms after (D) the onset of the synaptic stimulus. **E–G**: No LTD was induced by the stimulation protocol (1 Hz paired with post-synaptic stimulation for 2 min) in the absence of M1-receptor activation, but pairing-interval-dependent LTP was induced in presence of *β*-adrenergic inputs. **H–J**: Pairing-interval-dependent LTD was induced when the synaptic input was coupled with cholinergic inputs, and STDP was induced when both cholinergic and *β*-adrenergic inputs were present. **E, H**: Relative concentration of GluR1 at the membrane 16 min after the stimulation onset (normalised by concentration of membrane-inserted GluR1 at rest). **F, I**: Relative concentration of GluR2 at the membrane 16 min after the stimulation onset (normalised by concentration of membrane-inserted GluR2 at rest). **G, J**: Relative synaptic conductance (Eq. 5) 16 min after the stimulation onset (normalised by synaptic conductance at rest).

We first confirmed that the membrane expression of the glutamate receptors was not strongly affected by paired synaptic and somatic stimulation in the absence of *β*-adrenergic (which activates the PKA pathway) and cholinergic (which activates the PKC pathway) neuromodulation. Our model predicted that there is little change in the membrane expression of GluR1 and GluR2 type receptor subunits in this stimulation protocol (Fig. 6E–F, purple dashed line). Consequently, our model reproduced the observation [Seol et al., 2007] that this stimulation protocol led to little change in predicted synaptic conductance (Fig. 6G, purple dashed).

We next considered the paired synaptic-somatic stimulation in the presence of *β*-adrenergic ligand. Our model predicted a prominent (up to 75%) increase in GluR1 membrane expression with little effect on GluR2 membrane expression (Fig. 6E–F, blue). The predicted increase in GluR1 membrane expression (Fig. 6E) and the consequent increase in synaptic conductance (Fig. 6G, blue) were most prominent when the ISI was close to zero, and modest for large ISIs. These predictions are consistent with the experiments where an ISI-dependent potentiation of the EPSCs in the presence of *β*-adrenergic receptor agonists and absence of cholinergic agonists was observed [Seol et al., 2007].

When *β*-adrenergic neurotransmission was blocked but M1 receptors were activated, the model predicted a prominent (up to 65%) decrease in GluR2 membrane expression, with little effect on GluR1 membrane expression (Fig. 6H–I, purple dashed). Our model of synaptic conductance (Eq. 5) predicted a decrease in total conductance in a GluR1-GluR2-balanced synapse for this condition (Fig. 6J, purple dashed), which is in line with the experimental data [Seol et al., 2007]. The depression takes place throughout the tested ISIs, but the effect was smallest for ISIs very close to zero due to the counteracting effects of GluR1 membrane-insertion (Fig. 6H, purple dashed). Finally, when both *β*-adrenergic and cholinergic neurotransmission were active, our model predicted an increased GluR1 membrane expression and decreased GluR2 membrane expression, both of which were ISI dependent (Fig. 6H–I, blue). In these simulations, the predicted synaptic conductance was increased when the stimuli were coincident or near-coincident and decreased otherwise (Fig. 6J, blue), which is qualitatively similar to experimental data [Seol et al., 2007]. These results are dependent on the availability of both GluR1 and GluR2 subunits at the post-synaptic spine: in simulations where GluR2 or GluR1 subunits were absent, only LTP (extended data set, Fig. 4.1C) or LTD (Fig. 4.1D), respectively, was induced by the STDP protocol. In a similar manner as the HFS- and LFS-induced plasticity in Fig. 4.2A, we could reproduce the STDP using the dimer-of-like-dimers tetramer formation rule with a GluR1 fraction of 35% (Fig. 4.2B). Taken together, our model with balanced numbers of GluR1 and GluR2 subunits reproduces the neuromodulator-gated STDP observed in layer 2/3 pyramidal cells of the visual cortex.

### 3.5 The model predicts multimodal, protein concentration- and neuromodulation-dependent rules of plasticity

Cortical neurons express a variety of forms of LTP/LTD depending on the brain region and cell type. In computational studies, neocortical plasticity is most typically described by simple rules according to which small-amplitude Ca^2+^ inputs lead to depression of the synapse whereas large-amplitude inputs lead to potentiation. Apart from a few examples [Castellani et al., 2001, d’Alcantara et al., 2003, Castellani et al., 2005, Honda et al., 2013], these models typically do not describe the intracellular signalling machinery leading to the resulting plasticity [Holthoff et al., 2002, Karmarkar et al., 2002, Badoual et al., 2006, Cornelisse et al., 2007, Kubota and Kitajima, 2008, Urakubo et al., 2008]. Unlike biochemically detailed models, the simple models cannot be used to explore whether and how the prevalence of different plasticity-related proteins gives rise to various types of LTP/LTD or their impairments, which is an important question in the study of mental disorders with deficits in cortical plasticity. Here, we analysed the biochemical underpinnings of different types of plasticity rules using our unified model of cortical plasticity in order to predict the conditions for different forms of plasticity.

In a similar fashion to Section 3.2, we simulated our model of the post-synaptic spine when stimulated with a prolonged (5 min) square-pulse influx of Ca^2+^ and neuromodulators. We randomly altered the model parameters controlling the initial concentrations of different proteins, namely, the ratio of GluR1 to GluR2 subunits, the concentration of NCX (regulating the rate of Ca^2+^ decay from the spine), and the concentrations of PKA-pathway and PKC-pathway proteins (upstream of PKA and PKC). Alterations of the initial concentration of CaMKII (the only molecule in our model that exclusively affects the CaMKII pathway) had little effect in most domains of plasticity considered here (not shown), and thus, we omitted it in this analysis. We sampled these parameters from the following intervals: GluR1 ratio from the interval from 0 to 1 (keeping the total concentration of GluR subunits fixed at 540 nM), NCX concentration from the interval from 0 to twice the original value (2*×*0.54 mM), and the PKA and PKC-pathway factors *f*_PKA_ and *f*_PKC_ from the interval from 0 to 2 (see Section 2.4). We simulated the post-synaptic spine 75,000 times using different random values for these parameters under zero, low (50 particles/ms), medium (150 particles/ms), and high (250 particles/ms) levels of Ca^2+^ input.

We classified the parameter sets based on the total synaptic conductance 15 min after the onset of the stimulation with the high Ca^2+^ flux (250 particles/ms): the relative synaptic conductance varied between 0.16 and 5.92, and thus, we grouped the parameter sets to 16 classes using a bin size of 0.36 (Fig. 7A). We then analysed the parameter distributions and their co-variations within these classes and how the different parameters affected the shape of the LTP/LTD curve within each class, in particular, how strong LTP/LTD was induced by the medium (150 particles/ms) Ca^2+^ flux and whether the plasticity outcome was monotonic (in LTP, larger Ca^2+^ flux among the tested amplitudes always caused larger synaptic conductance; in LTD, larger Ca^2+^ flux always caused smaller synaptic conductance) or not. A special subset of the non-monotonic forms of LTP were the BCM-type plasticity curves, where either 50 or 150 particles/ms Ca^2+^ injection resulted in LTD and the 250 particles/ms resulted in LTP.

**Figure 7:**
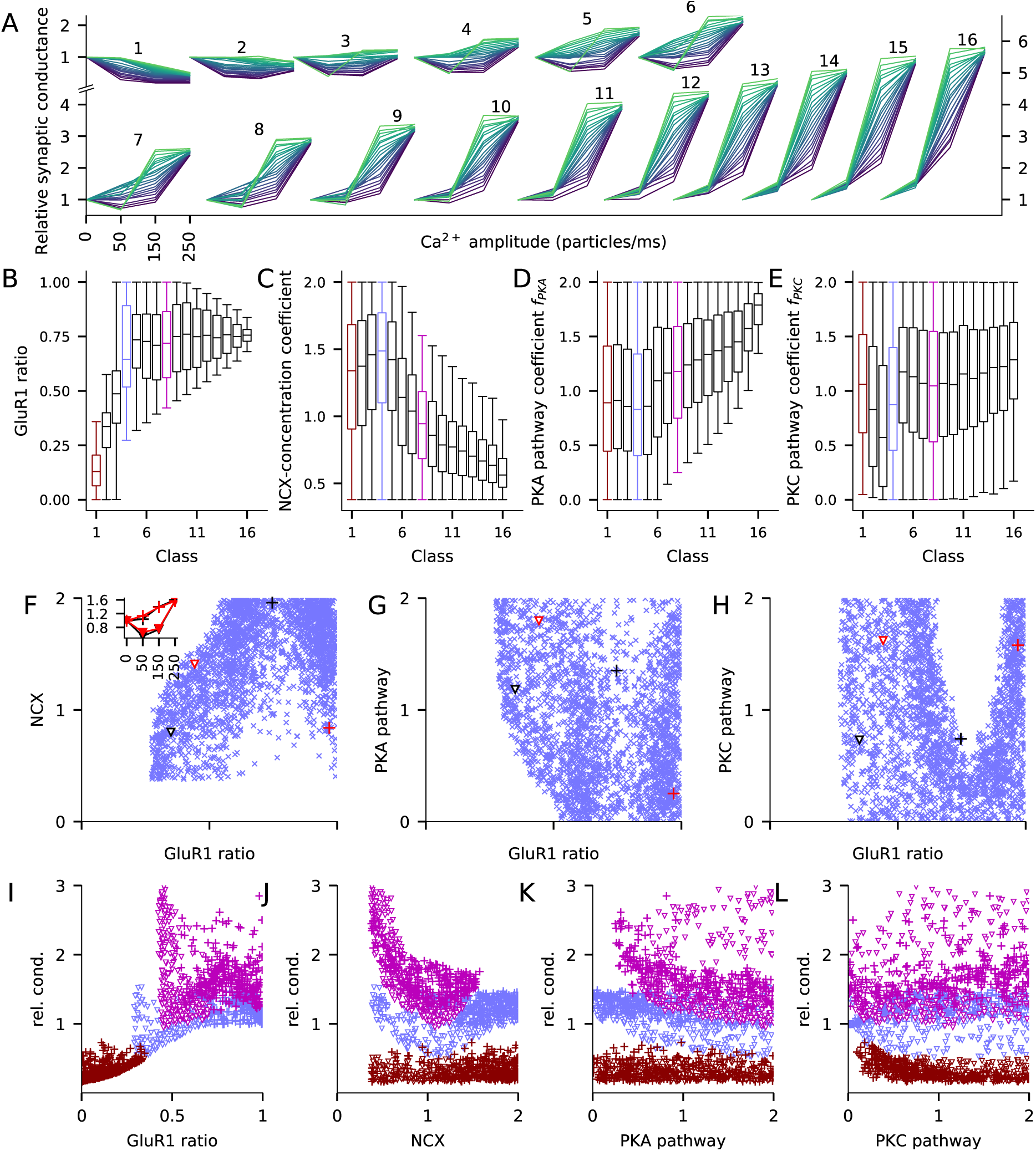
The fraction of GluR1s, number of Ca*^2+^* expulsion proteins, and the concentrations of PKA and PKC-pathway proteins in the post-synaptic spine determine the type of LTP/LTD in the post-synaptic spine. A: The LTP/LTD curves are displayed for all 16 classes. Four values of Ca^2+^ input amplitude were considered: 0, 50, 150, and 250 particles/ms (x-axis; repeated and overlaid for space). The y-axis shows the relative synaptic conductance, i.e., total synaptic conductance 15 min after the onset of the Ca^2+^ input divided by the total synaptic conductance before the Ca^2+^ input. 20 representative parameter sets are displayed from each class, coloured from purple (lowest relative synaptic conductance response for medium Ca^2+^ input) to green (highest conductance). **B–E**: Distribution of model parameters, i.e., GluR1 ratio (B), NCX-concentration coefficient (C), PKA pathway-concentration coefficient *f*_PKA_ (D), and PKC pathway-concentration coefficient *f*_PKC_ in the 16 classes. Classes 1 (brown), 4 (blue), and 8 (magenta) highlighted for further analysis. **F–H**: GluR1 ratio plotted against NCX-concentration coefficient (F), *f*_PKA_ (G), and *f*_PKC_ (H) in class 4. The red and black markers pinpoint two pairs of parameter sets, two of them (triangles) producing similar non-monotonic, BCM-type plasticity and two of them similar monotonic LTP curve (inset of F; see panel A). **I–L**: Relative conductance in response to medium Ca^2+^ input plotted against GluR1 ratio (L), NCX-concentration coefficient (M), *f*_PKA_ (N), and *f*_PKC_ (O) in classes 1, 4, and 8. ‘+’-markers represent parameter sets that induced monotonic LTP/LTD curve (within the data for Ca^2+^ flux of 0, 50, 150, and 250 particles/ms), and triangles represent non-monotonic LTP/LTD.

Classes 11–16 exhibited the strongest LTP, with large synaptic conductance for both 150 and 250 particles/ms Ca^2+^ injection, whereas classes 1 and 2 only exhibited LTD (Fig. 7A). Classes 3–12 exhibited BCM-type of plasticity but the majority of the LTP/LTD curves were of non-BCM type in each class (Fig. 7A).

Three parameters — the GluR1 ratio, NCX concentration and *f*_PKA_, differed significantly across the 16 classes (Fig. 7B–D). Low GluR1 ratio was needed for strong LTD and medium or high GluR1 ratio for strong LTP (Fig. 7B). However, the strongest forms of LTP (classes 11-16) were induced only when GluR1 ratio was smaller than 1 (Fig. 7B), because a very low number of GluR2 subunits implied that the synapse has many homomeric GluR1 tetramers at a basal state, and thus stimulation-induced GluR1 exocytosis and GluR2 endocytosis did not radically increase the number of homomeric GluR1 tetramers (Eq. 5, see also Fig. 5D). For LTD and moderate LTP (classes 1–5), any NCX concentration and PKA-pathway coefficient could be used, but very strong LTP (classes 10-16) required a small to medium NCX concentration (Fig. 7C) and a medium to large PKA-pathway coefficient (Fig. 7D). By contrast, PKC-pathway coefficient alone was not predictive of plasticity outcome (Fig. 7E).

The model results for the large parameter distributions of Fig. 7B–E imply that there are manifestly different combinations of parameters that lead to the same LTP/LTD outcome. To analyse this intrinsic variability, we studied the distributions of the model parameters within the class of mild LTP (class 4, 24–60% LTP for 250 particles/ms; indicated by blue boxes in Fig. 7B–E) in more detail. Class 4 spanned second-largest range of GluR1-ratio (values from 0.28 to 1). Dependencies between the four parameters could be observed in 2-dimensional plots of the parameter space (Fig. 7F–H). A large GluR1 ratio was typically accompanied by a large NCX concentration, except for the highest (*>* 0.75) GluR1 ratios (Fig. 7F). Furthermore, large PKA-pathway coefficient *f*_PKA_ was required in this LTP class if GluR1 ratio was small (Fig. 7G). As for the contribution of PKC-pathway coefficient, for medium to large *f*_PKC_, the GluR1 ratio was bimodally distributed, while for small *f*_PKC_, any GluR1 ratio within the range 0.28 to 1 could produce class-4 plasticity (Fig. 7H). Typically, we could find many parameter sets producing similar LTP/LTD curves: the red and black data points illustrate two pairs of parameter sets that are different from each other in all four parameters (Fig. 7F–H) but produced same types of monotonic (’+’ markers) or BCM-type (triangles) plasticity curves (Fig. 7F inset).

In addition to the intra-group diversity in the above four parameters (Fig. 7B–E), there was a large variability in the shape of the LTP/LTD curves within each class despite the strictly constrained LTP/LTD outcome for large Ca^2+^ inputs (Fig. 7A). To analyse the effects of the model parameters on the LTP/LTD curve shape, we determined the correlation coefficients between the four parameters and the relative synaptic conductance in response to the medium Ca^2+^ input (150 particles/ms) in three of the above classes. GluR1 ratio (Fig. 7I) was positively correlated with the synaptic conductance in response to medium Ca^2+^ input in both classes of strong LTD (class 1, 48–84% LTD, brown; correlation coefficient 0.56 — n.b., this is an anticorrelation with the magnitude of the LTD) and moderate LTP (class 4, blue; correlation coefficient 0.58), but not in the class of strong LTP (class 8, 168–204% LTP, magenta; correlation coefficient −0.09). This correlation was strong in the class of non-monotonic LTD (class 1; correlation coefficient 0.80 among the parameter sets inducing non-monotonic LTD curve and 0.50 among those inducing monotonic LTD). The lack of correlation in the class of strong LTP is likely due to a large basal synaptic conductance: if absolute instead of relative synaptic conductance were used, the correlation coefficient was 0.67 in class 8. The synaptic conductance in response to medium Ca^2+^ input was also moderately anticorrelated with NCX concentration in class 8 (correlation coefficient −0.62, Fig. 7J) and with PKC-pathway coefficient in class 1 (correlation coefficient −0.52, Fig. 7L). The former result can be explained by the fact that an increase in NCX concentration decreases the Ca^2+^ transients, thus decreasing the activation of PKA and CaMKII pathways, and the latter result is consistent with our observations on the effects of PKC-pathway-activating ligands on the magnitude of LTD (Fig. 5E–G). The PKA-pathway coefficient was to some degree negatively correlated with the synaptic conductance in response to medium Ca^2+^ input (correlation coefficient was −0.09, −0.48, and −0.35 in classes 1, 4, and 8, respectively; Fig. 7K). Among the class-8 parameters inducing monotonic LTP, the synaptic conductance in response to medium Ca^2+^ input was anticorrelated with the PKA-pathway coefficient (correlation coefficient −0.62, Fig. 7K) — this is consistent with the correlation between NCX and PKA-pathway coefficient within this subclass (not shown) and the aforementioned anticorrelation of NCX and synaptic conductance in response to medium Ca^2+^ input (Fig. 7J).

Taken together, our results show that alterations of the concentrations of the proteins regulating Ca^2+^ efflux or PKA/PKC-pathway signalling and the numbers of GluR1 and GluR2 subunits, ranging from complete absence to moderate increase (*±*100%), have a large effect both on the type of plasticity (LTP or LTD) that the synapse can express and on the sensitivity of the plasticity outcome to the amplitude of the Ca^2+^ flux.

### 3.6 A parametric analysis confirms the robustness of the model

We analysed the model responses to 4xHFS and LFS protocols (as in Fig. 4) under small (*±*10%) changes in the parameters describing the initial concentrations and reaction rates (Fig. 8). As expected, most parameter changes led to small deviations from the predicted magnitudes of LTP/LTD (Fig. 8, grey bars). Alterations of the initial concentration of a number of species (7 out of 47) and reaction rates (13 out of 413) resulted in a notable (*>*15%) amplification or attenuation of LTD (Fig. 8A) or LTP (Fig. 8B). The parameters affecting the LFS-induced LTD were all related to GluR1 membrane insertion or total amount of GluR1 (Fig. 8A), while the parameters affecting the 4xHFS-induced LTP were related to NCX-mediated Ca^2+^ expulsion, PP1 concentration, production of cAMP by AC1, PKA buffering/deactivation, or GluR1 membrane insertion (Fig. 8B). Importantly, none of the parameter changes completely abolished the LTP or LTD. Taken together, our model is robust to small alterations in initial concentrations and reaction rates, but parameters influencing the Ca^2+^ dynamics, GluR1 activity, or the PKA-pathway signalling can have relatively large effects on the model output.

**Figure 8:**
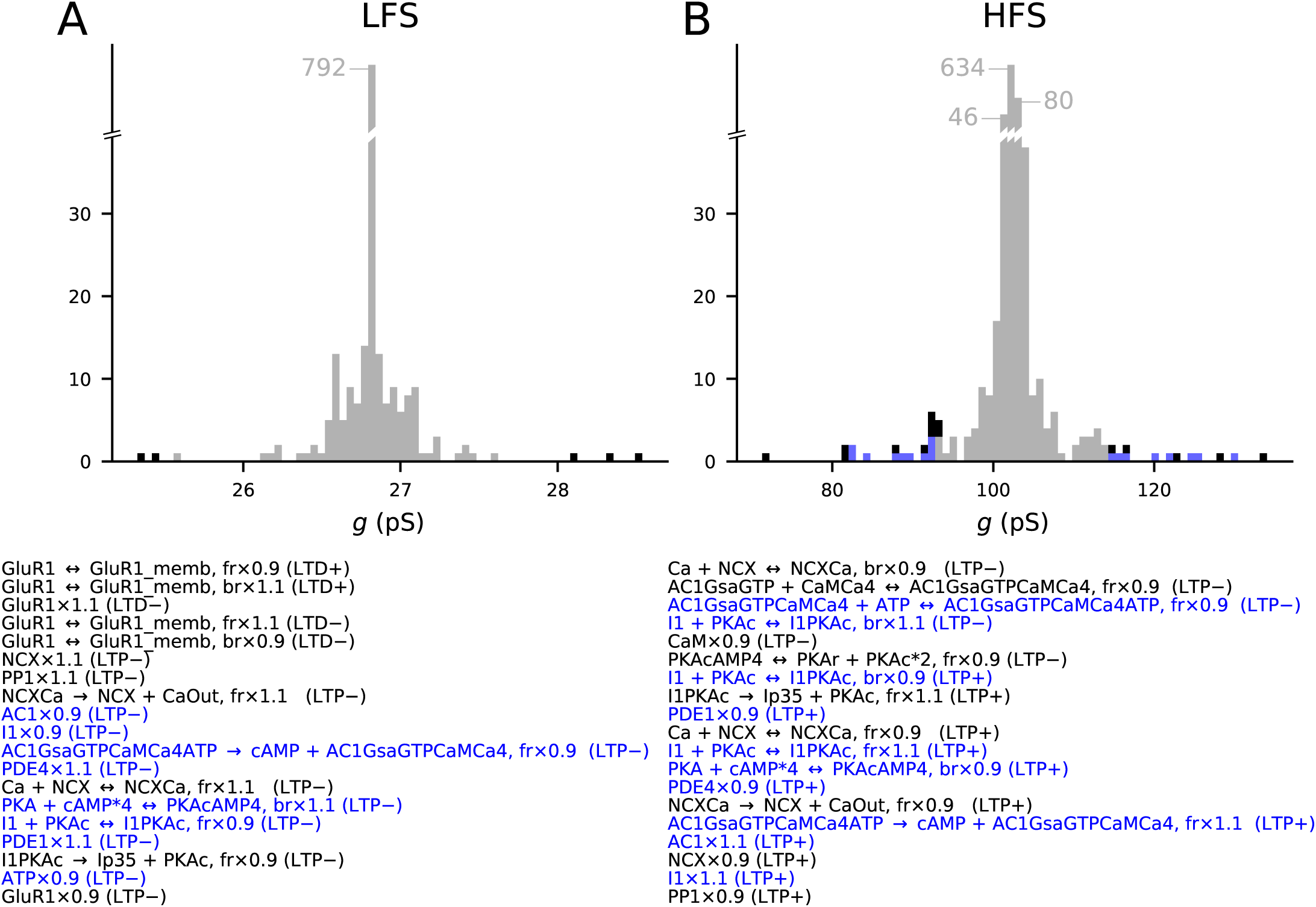
The model predictions of LTP and LTD are robust to small changes in model parameters. Values of initial concentrations (47 parameters) or reaction rates (413 parameters) were changed one at the time by −10% or +10%, and the resulting synaptic conductance 16 min after LFS (A) or 4xHFS (B) protocol was measured (NEURON RxD simulations). The initial synaptic conductance is 33.4 pS (see Fig. 4A,S), although some parameter changes mildly affected this value (data not shown). The x-axis shows the post-LFS (A) or post-HFS (B) synaptic conductance, and the y-axis shows the number of parameter alterations. *±*10% changes in initial concentrations of 9 species and 11 reaction rates caused *>* 15% change in the amplitude of LTP or LTD — these changes are represented by black (multi-pathway parameters) and blue (PKA-pathway parameters) bars. The underlying parameter changes are listed below the figure in the order they appear in panels A and B: first the strengthened LTD (left-hand tail of A), followed by weakened LTD, weakened LTP, and finally strengthened LTP (right-hand tail of B).

### 3.7 The model flexibly reproduces data from various cortical LTP/LTD experiments

The richness of the intracellular signalling machinery behind LTP and LTD poses challenges for both qualitative and quantitative comparison between results from different cell types, obtained using different stimulation protocols, or even published by different laboratories [Larkman and Jack, 1995]. Computational biochemically detailed models have been proposed as an absolutely reproducible tool that is particularly suited for unifying our understanding of LTP and LTD across cell types and brain regions [Manninen et al., 2010]. Here, we show that our model for intracellular signalling in a cortical post-synaptic spine — through the use of varying concentrations of different proteins — can be flexibly tuned to reproduce data from the experimental literature of cortical LTP/LTD. This allows one to make predictions for the differences in intracellular machineries underlying each of the experiments, leading to a more complete view of the plasticity-related signalling pathways in different cell types in the cortex and the effects of the stimulation protocol on the plasticity outcome across cortical areas.

To show the flexibility of our model, we aimed to reproduce a large amount of data on cortical plasticity across cortical areas and stimulation paradigms. We reviewed the literature of cortical plasticity, and picked 8 studies where one or more types of neurons were tested using one or more stimulation protocols and the outcome was quantified using electrophysiological measurements (Tab. 4). These studies comprised 11 data sets that described the response of a neuron population in entorhinal cortex (EC), prefrontal cortex (PFC), barrel cortex (BC), anterior cingulate cortex (ACC), visual cortex (VC), or a culture from auditory cortex (AC) to plasticity inducing protocols (Tab. 4). For each experiment in each data set, we assigned an objective function that quantified the error between the predicted LTP/LTD outcome (measured in relative synaptic conductance) and the data (typically measured in fold change of field EPSP slope). The objective functions were averaged across different time instants (10, 15, and 20 min post-stimulus-onset). We then ran a multi-objective optimisation algorithm (see Section 2.4) that aimed at finding the values for model parameters that minimised these objective functions. The fitted parameters included the amplitudes of pre-synaptic stimulation-associated fluxes of Ca^2+^, *β*-adrenergic ligand and glutamate in addition to GluR1 fraction and factors for the protein concentrations of different pathways. We ran the optimiser for 20 generations. For data sets VC-1 and VC-2 we did not find parameter sets that would fulfil all four objective functions, and therefore, we re-fitted the model for these data sets excluding the CaMKII-blocked experiments (printed in grey in Tab. 4). We found groups of parameter sets that fit within one standard deviation (SD) on average from the target values of synaptic conductance for each data set of Tab. 4 (Fig. 9A–C). There were differences in the numbers of acceptable parameter sets between the data sets due to differences in the postulated strength of the LTP/LTD, the number of experiments, and the SD of the post-stimulus synaptic conductance (Fig. 9D). The data sets EC-1 and BC were particularly challenging to fit (*<*0.2% of the parameter sets tested by the optimiser gave an acceptable fit; Fig. 9D). By contrast, the data set AC-2 was the easiest to fit (3.9% of the parameter sets were acceptable; Fig. 9D).

**Figure 9:**
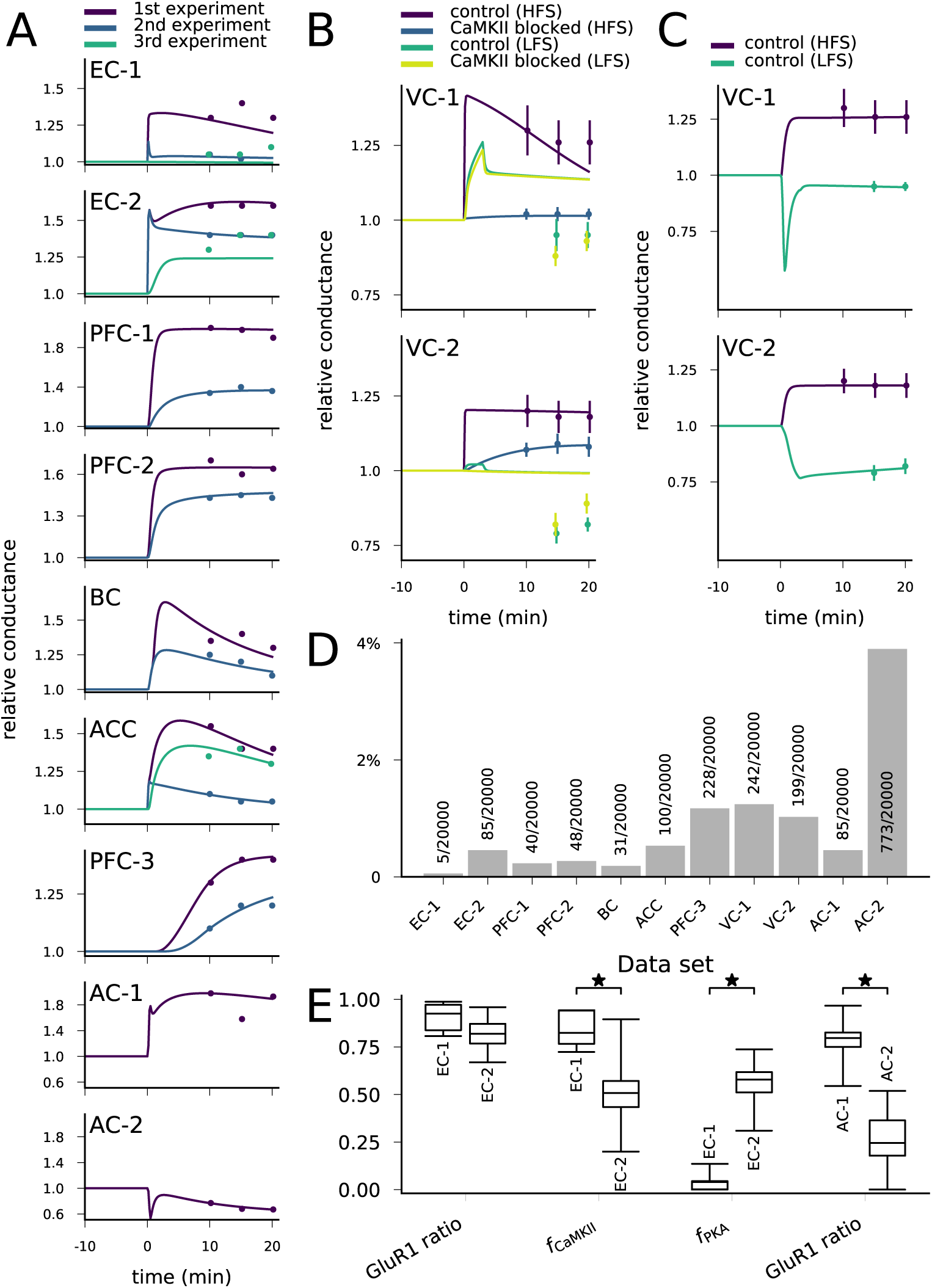
**The model can be fit to LTP/LTD data from different cortical areas**. **A**: The model could be fit to LTP/LTD data from data sets EC-1 (top), EC-2, PFC-1, PFC-2, BC, ACC, PFC-3, AC-1, and AC-2 (bottom). The curves represent the model predictions of the best-fit parameter sets, and the dots represent the experimental data from Tab. 4. For data sets other than AC-1 and AC-2, several experiments with various chemical agents or genetic mutations were performed for each neuron population: these are ordered as in Tab. 4 (e.g., in data set EC-1, purple (1st experiment) corresponds to control, blue (2nd experiment) to CaMKII-blocked experiment, and green (3rd experiment) to the experiment where post-synaptic Ca^2+^ was blocked). **B**: The model could not be fit to the complete LTP/LTD data from data sets VC-1 (top) and VC-2 (bottom). The best parameter sets correctly predicted the LTP/LTD in up to two experiments (e.g., the selected parameter sets reproduce the HFS data with and without CaMKII inhibitor, but failed to reproduce the LFS data). **C**: The model could be fit to the LTP/LTD data from data sets VC-1 (top) and VC-2 (bottom) when CaMKII-blocked experiments were ignored. The vertical bars in (B) and (C) represent the SD from the experimental data. **D**: Proportion of accepted parameter sets across the 20 generations of multi-objective optimisation (20’000 parameter sets in total) in each data set. **E**: Box plots of selected parameters in the acceptable parameter sets of data sets EC-1 and EC-2 (three left-most pairs) and AC-1 and AC-2 (right-most pair). Values of *f*_CaMKII_ and *f*_PKA_ are linearly scaled such that the values 0 and 1 correspond to 0 and double the original value of the underlying parameters, respectively (CaM and CaMKII for *f*_CaMKII_, and R, Gs, AC1, and AC8 for *f*_PKA_, see Section 2.4). The medians were significantly different in the compared data sets (U-test, p-value *<* 0.001).

The obtained parameters reflect the pathways needed for the type of plasticity. For example, the LTP of synapses of the horizontal but not those of the ascending pathway to EC were blocked by CaMKII inhibition, while the LTP of synapses of the ascending pathway were blocked by PKA inhibition [Ma et al., 2008]. This is reflected in the obtained parameter sets: the parameter controlling CaM and CaMKII concentrations (*f*_CaMKII_) was significantly larger (U-test, p-value *<* 0.001) in the horizontal-pathway (data set EC-1) synapse models, while the parameter controlling upstream PKA-pathway proteins (*f*_PKA_) was significantly larger in the models reproducing the data from the ascending pathway (data set EC-2; Fig. 9E). The GluR1-GluR2 ratio, in turn, was not significantly different between the two data sets (Fig. 9E). As a contrasting example, our model predicts a large variety of parameters that reproduce the LTP and LTD of data sets AC-1 and AC-2 but, consistent with the results of Fig. 7, the GluR1 ratio was significantly larger in the model parameters fitted to the data from LTP-expressing neurons than from LTD-expressing neurons (Fig. 9E). The complete graphs of parameter value distributions in the 11 data sets and the parameter set producing the best fit (Fig. 9A) for each data set are shown in the extended data set (Figs. 9.1–9.11). Taken together, our model-fitting experiment shows that the model can be fit to many types of multi-condition plasticity data — without altering the reaction rates — and that the resulting predictions of the underlying protein concentrations reflect the mechanisms proposed by the experimental studies.

The models obtained by fitting the initial concentrations to data provide an important tool for predicting the outcome of plasticity under various stimulus protocols and chemical agents. We carried out additional simulations with the obtained models using HFS protocol. The models fitted for data from EC (data sets EC-1 and EC-2, [Ma et al., 2008]), BC [Hardingham et al., 2003], ACC [Song et al., 2017], and LTP-expressing cultured cortical neurons (data set AC-1, [Kotak et al., 2007]) predicted a steady increase in response to HFS, while models fitted for other cortical data predicted a mixture of LTP, LTD, and no change (Fig. 10A). Furthermore, to obtain experimentally testable predictions for the dependency of the plasticity outcome in different cortical areas on the intracellular signalling, we simulated the models from each data set with the corresponding stimulus protocol with CaMKII, PKA, or PKC blockade. The inhibition of CaMKII impaired the LTP in data set EC-1 (horizontal pathway) but had little or no effect on the plasticity in other cortical areas (Fig. 10B). The lack of effect of CaMKII blockade on the ascending pathway of EC — an experiment which was not included in the fitting of the model (Tab. 4) — validates the underlying models in this aspect since similar results were observed in [Ma et al., 2008]. Moreover, the similarities in the predicted effect of CaMKII-, PKA-, and PKC-pathway blockades between the two models of CC*→*PFC synapses (PFC-1 and PFC-2; Fig. 10B–D) serve as an additional validation of these models. The inhibition of PKA impaired LTP in all cortical areas, except for LTP in the horizontal pathway of the EC (EC-1; Fig. 10C). Our models also predicted that LTP of the CC*→*PFC pathway and LFS-induced LTP in VC can be effectively weakened or even be transformed to a mild LTD by PKA blockade (Fig. 10C). Finally, our models predicted that PKC inhibition transformed all forms of LTD (LFS-induced LTD in VC-1 and VC-2; LTD in cultured cortical neurons, AC-2) into LTP and impaired certain forms of LTP (LTP in EC-2; LTP in cultured cortical neurons, AC-1) (Fig. 10D). Taken together, our results suggest that almost all forms of post-synaptic plasticity in the cortex are likely to be PKA-dependent, and that many types of cortical plasticity are also influenced by CaMKII and PKC activity. Our results highlight the need for additional chemical or genetic manipulations to be done when experimenting on cortical plasticity in order to correctly reveal the intracellular signalling cascades in the post-synaptic spine.

**Figure 10:**
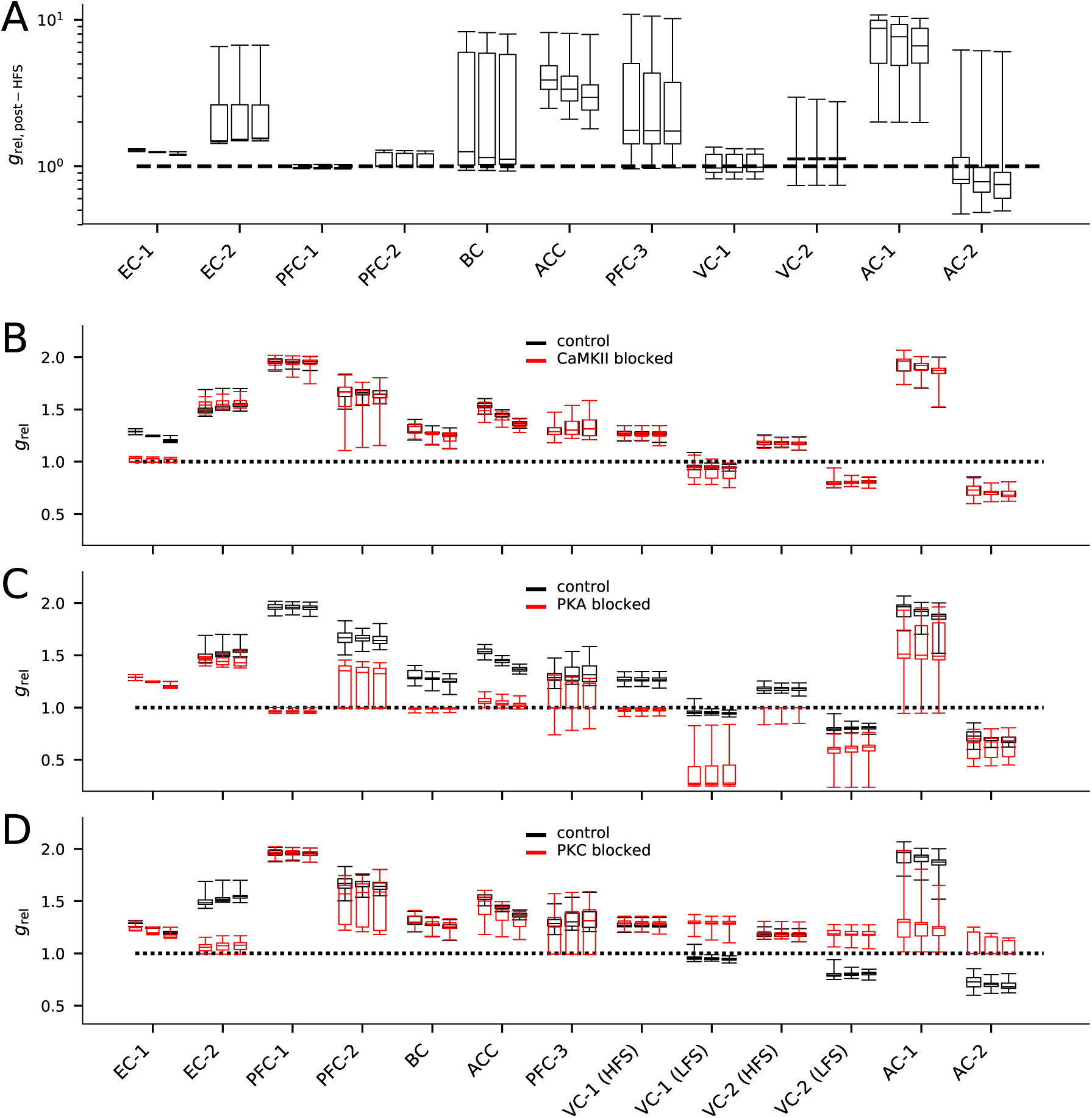
The models describing plasticity in different cortical areas predict diverse responses to modified stimulation protocol and stimulation under chemical blockers. A: The predicted responses of the 20 best models in each data set to HFS (100 pulses at 100 Hz) stimulation. **B–D:** The predicted responses of the 20 best models in each data set to the applied stimulation protocol (see Tab. 4) when CaMKII (B), PKA (C), or PKC (D) activity was blocked (red) or under control condition (black).

## 4 Discussion

We built a single-compartment model describing the major post-synaptic signalling pathways leading to LTP and LTD in the cortex. We showed that our model reproduced conventional types of LTP and LTD, where an HFS-induced increase in GluR1 can increase the synaptic conductance (LTP) and an LFS-induced endocytosis of GluR2 can decrease it (LTD; Fig. 4) and reproduced STDP data from visual cortical layer 2/3 pyramidal cells (Fig. 6). Our model explains how different forms of plasticity depend on the concentrations of PKA- and PKC-pathway proteins (Fig. 7). We also showed that our model can be fit to explain the pathway dependencies of various types of neocortical LTP/LTD data published in the literature (Fig. 9). Our fitted models provide a powerful tool for testing hypotheses on the effects of chemical or genetic manipulations on the LTP and LTD in different cortical regions (Fig. 10).

### 4.1 Role of GluR2 in synaptic plasticity in the neocortex

GluR2 subunits are highly expressed in neocortical neurons [Kondo et al., 1997], and their endocytosis mediates (or, at minimum, is correlated with) synaptic depression in many cortical regions such as ACC [Toyoda et al., 2007], VC [Heynen et al., 2003], and PFC [Van den Oever et al., 2008]. Previous intracellular signalling-based models of neocortical LTP/LTD exist, but they do not take into account the contributions of GluR2 subunit. For example, in three previous models [d’Alcantara et al., 2003], [Castellani et al., 2005], and [Honda et al., 2013], S831-phosphorylation-mediated LTP was described in a fashion similar to our model (although more approximations were made), but the phosphorylation site S845 was assumed to be basally phosphorylated and LTD was caused by modest Ca^2+^ inputs that led to PP1 or PP2B-mediated dephosphorylation of S845. Although there is support for this order of events [Lee and Kirkwood, 2011, Diering et al., 2016], newer findings have confirmed the low degrees of phosphorylation of both S831 and S845 at a basal state in cortical cells, especially in synaptic spines [Diering et al., 2016]. To analyse the contributions of GluR2 subunits to neocortical LTP/LTD, we included in our model the signalling pathways leading to both phosphorylation-mediated exocytosis of GluR1 and endocytosis of GluR2. Our model could thus be used to study not only PKA-mediated LTP or PKC-mediated LTD but also their co-effects and co-dependencies.

### 4.2 Implications of the study

The modelling results of the present work give rise to experimentally testable predictions. For example, our STDP model, when stimulated without *β*-adrenergic ligands, suggests that at near-zero pairing-intervals the magnitude of the depression may be decreased or even switched to mild LTP (Fig. 6J). In many experimental studies (including [Seol et al., 2007]), the type and magnitude of plasticity in this regime of STDP is not reported. We also predict that a mild LTP (24–60% LTP; class 4 in Fig. 7) can be obtained through many differently weighted interactions of PKA and PKC pathways and Ca^2+^ expulsion strengths (Fig. 7F–H, 9.1-9.2, 9.5-9.7). Importantly, this is the regime of a wealth of experimental LTP data ([Tsumoto, 1990]; cf. Tab. 4), which is consistent with the great diversity of LTP mechanisms observed in the neocortex [Feldman, 2009]. Based on our simulated data (Fig. 10), we suggest that in order to correctly characterise the mechanisms behind LTP of especially this magnitude, both experiments that activate the PKA pathway and experiments that block or activate the PKC pathway should be carried out.

A key challenge in the study of synaptic plasticity is the diversity of LTP/LTD observed across the cell types in the brain [Granger and Nicoll, 2014]. Differences in the transcriptome has been proposed as one of the sources for this variability (cf. [Lisachev and Shtark, 2018]). We believe our model can be used to explain some of the discrepancies in the experimental data in this regard and expand the understanding of possible molecular contributors to LTP/LTD. For example, it is known that activation of PKA pathway by dopamine or noradrenaline in PFC pyramidal neurons increases the synaptic conductance through GluR1 membrane insertion [Sun et al., 2005, Xu et al., 2010]. Our model is in agreement with this (Fig. 4), but it also proposes that the LTP can be impaired by over- or underexpression of many involved proteins, such as AC1, I-1 (inhibitor of PP1), PDE4, PDE1, GluR1, and CaM, and even alterations in ATP concentration (see Fig. 8B). Small differences in the concentrations of a number of such contributing proteins are likely to cause significant alterations to LTP observed in different brain areas and cell types.

### 4.3 Validity of the results and limitations of the study

Our model of total synaptic conductance of the post-synaptic spine is based upon a number of assumptions. First, the prediction of a large increase of conductance that follows the replacement of GluR2 subunits at the membrane by GluR1 subunits (e.g., Fig. 4) is based upon the findings on differences in single-channel conductances of different types of AMPAR tetramers in hippocampal neurons [Oh and Derkach, 2005]. Following [Oh and Derkach, 2005], we assumed that CaMKII-phosphorylation of S831 only increases the conductance of GluR1 homomers and not that of GluR1/GluR2 heteromers, although also heteromers have been observed to increase their conductance in the presence of transmembrane AMPAR regulatory proteins [Kristensen et al., 2011]. Second, we assumed a random tetramerization procedure in which each of the four subunits in the tetramer may be either GluR1 or GluR2 subunit. Traditionally, AMPARs were thought to assemble as dimers of like dimers, i.e., that first GluR1s and GluR2s assemble into homomeric R1-R1 and R2-R2 dimers and R1-R2 heterodimers and that these three types of dimers only assemble into tetramers with a dimer of its own kind [Gan et al., 2015]. However, recent findings of heterotetramers with only one GluR1 subunit [Zhao et al., 2019] challenge this model. To show that our results were not dependent on the details of this process, we reproduced our results using the alternative (dimer of like dimers) tetramer formation rule. Using a slightly modified GluR1-GluR2 balance (35:65), this model reproduced HFS-induced LTP and LFS-induced LTD (Fig. 4.2A) as well as the neuromodulator-gated STDP (Fig. 4.2B). In summary, our model predictions were not dependent on the assumptions on the tetramer formation rule.

Our model reproduced the qualitative results of STDP of layer 2/3 pyramidal cells in visual cortex being gated by neuromodulators, but there were quantitative differences. When acetylcholine was present, our model predicted a prominent decrease in GluR2 membrane-expression regardless the pairing interval (Fig. 6I), which caused a notable LTD for very large pairing intervals (Fig. 6J), whereas the experimental data showed attenuation of the depression for large inter-stimulus intervals [Seol et al., 2007]. This discrepancy is likely caused by slower time-scale (*>* 50 ms) processes that are either not included (e.g., Ca^2+^-induced Ca^2+^ release or cAMP-dependence of HCN channels) or not adequately strong (e.g., contributions of voltage-gated Ca^2+^ channels or SK channels, cf. [Mäki-Marttunen et al., 2017]) in the multi-compartmental model. On the other hand, mechanisms lacking from the biochemical model (e.g., voltage-dependence of the Ca^2+^-expulsion rate of NCX [Weber et al., 2002]) could also impede our results. Some aspects of cellular physiology could therefore be better represented if we incorporated both biochemical signalling and multicompartmental Hodgkin-Huxley-type modelling into the simulations, as done in modelling studies of persistent neuron firing [Neymotin et al., 2016] and astrocyte electrophysiology [Savtchenko et al., 2018].

The quality of the model fitting to experimental data in Section 3.7 is restricted by the fact that not all of the LTP/LTD data in Tab. 4 were confirmed to have a post-synaptic origin. This may be the key source of discrepancy in the fitting of the model to the CaMKII-blocked data from [Kirkwood et al., 1997] (Fig. 9B), since CaMKII activation at the pre-synaptic spine may lead to EPSC potentiation through an increase in neurotransmitter release [Ninan and Arancio, 2004]. This scenario is supported by [Seol et al., 2007] where S831-deficient mice were observed to show normal post-synaptic LTP in the VC.

### 4.4 Outlook

Our results on interactions of the different pathways in post-synaptic spines including both GluR1 and GluR2 subunits provide valuable insights on the contributions of protein expression on the plasticity of the synapse. Previously, synaptic plasticity outcomes in the cortex have been conjectured to depend on the type of the post-synaptic cell type, in addition to the timing and frequency of the applied stimuli and dendritic filtering properties [Bi and Poo, 1998, Sjöström et al., 2001]. Our model provides a way to analyse exactly which aspects of PKA-, PKC- and CaMKII-pathway signalling underlie these cell-type-dependent differences in synaptic plasticity. Moreover, our model can be used for initial testing of hypotheses concerning dysfunctions (including chemical and genetic manipulations) of many intracellular signalling proteins and their role in impairments of cortical synaptic plasticity. By altering the initial concentrations or reaction rates of various species according to disease-associated functional genetics data, the model can be used to provide insights into the disease mechanisms of mental disorders that express both genetic disposition of post-synaptic signalling pathways and plasticity-related phenotypes, such as schizophrenia [Devor et al., 2017].

## 5 Acknowledgements

UNINETT Sigma2 resources (project NN9529K) were used for simulations. Funding: Research Council of Norway (248828), European Union Horizon 2020 Research and Innovation Programme under Grant Agreement No. 785907 [Human Brain Project (HBP) SGA2].

## 6 Financial disclosures

The authors declare that there are no conflicts of interest.

## 7 Author contributions

Designed the study: TMM, KTB

Provided data or analytical support: NI, AGE, GTE

Performed the analysis: TMM

Interpreted the results: TMM, NI, AGE, GTE, KTB

Wrote the manuscript: TMM

